# The First High-Quality Genome Assembly and Data Analysis of the Malaysian mahseer (*Tor tambroides*)

**DOI:** 10.1101/2022.01.02.474749

**Authors:** Melinda Mei Lin Lau, Leonard Whye Kit Lim, Hung Hui Chung, Han Ming Gan

## Abstract

The Malaysian mahseer (*Tor tambroides*), one of the most valuable freshwater fish in the world, is mainly targeted for human consumption. The mitogenomic data of this species is available to date, but the genomic information is still lacking. For the first time, we sequenced the whole genome of an adult fish on both Illumina and Nanopore platforms. The hybrid genome assembly had resulted in a sum of 1.5 Gb genomic sequence from the 44,726 contigs found with 44 kb N50 length and BUSCO genome completeness of 84.3%. Four types of SSRs had been detected and identified within the genome with a greater AT abundance than that of GC. Predicted protein sequences had been functionally annotated to public databases, namely GO, KEGG and COG. A maximum likelihood phylogenomic tree containing 53 Actinopterygii species and two outgroups was constructed, providing first insights into the genome-based evolutionary relationship of *T. tambroides* with other ray-finned fish. These data are crucial in facilitating the study of population genomics, species identification, morphological variations, and evolutionary biology, which are helpful in the conservation of this species.

## Introduction

The Malaysian mahseer, *Tor tambroides* (Bleeker, 1854), one of the members of the family Cyprinidae, is a widespread species found in aquaculture and fisheries mainly targeted for human consumption (Kottelat *et al*., 2018). It is commonly named Kelah or Empurau in Malaysia and Jurung, Indonesia (Jaafar et al., 2021). As a true mahseer (*Tor* spp.), it can be found in rapidly-flowing waters with rocky bottoms (Shreshtha, 1997). Together with *Tor tambra* and *Tor dourenensis*, it is one of the three *Tor* spp. found in freshwaters of Malaysia, and among the 16 *Tor* spp. found worldwide (Ng, 2004).

Like other *Tor* spp., *T. tambroides* is endangered by environmental degradation within their habitat, causing an elevating shrink in their population size in recent years (Ingram et al., 2005). Anthropogenic modification of rivers, including agricultural activities, logging, and deforestations, had interrupted and reduced the water flow within its habitat. Consequently, such actions not only shrink the population size of *T. tambroides* and other Tor spp., but it is also associated with impacts on the aquatic environment. Despite environmental issues, *T. tambroides* are also at threat from overfishing with the usage of hooks, nets, and dynamites. However, there is currently no information on the rate of these losses (Kottelat et al., 2018).

It resides a wide range of freshwaters, including Brunei Darussalam, Yunnan China, Indonesia (Kalimantan, Jawa, and Sumatera), Lao, Malaysia (Peninsular, Sabah, and Sarawak) and Thailand (Kottelat 2018). Despite its comprehensive habitat coverage, its distant population can probably be different, leaving its taxonomy still vague. Its body is covered with scales depending on locality, such as silver, bronze, and reddish.

Although the presence of an upper median projection can be observed in *T. tambroides*, its other features, including equal caudal fin lobe, long lower median lobe, sub-terminal mouth position and pointed rostrum hood, were shared closely with other *Tor* spp., making it difficult to be distinguished (Jaafar et al., 2021). Therefore, disentangling *Tor* spp. via lips polymorphisms and median lobes resulting from direct environmental influences is discouraged due to lack of direct evidence (Roberts & Khaironizam, 2008). Unpublished research had emphasised the reduction in lips and lobe size and the loss of red colour when the fishes from Sungai Tembat, Terengganu, were kept in captivity for more than two years (Walton et al., 2019). However, such polymorphism is yet to be observed in nature and thus remains deficient.

The accumulation of significant genetic and/or morphological differences among *Tor* spp distributed across Indonesia and Malaysia urged the revision of *Tor* taxonomy. Studies had been done aiming to resolve the ambiguous taxonomical status of *Tor* spp. by looking into its phylogenetic relationship through mitochondrial DNA, Cytochrome c Oxidase Subunit I (*COX1*) gene, Cytochrome b, ATPase 6 /8 gene, 16S rRNA gene, microsatellite, and SNP markers (Walton et al., 2017; Jaafar et al., 2021; Lim et al., 2021a). However, to date, the whole genome and transcriptome sequences of *Tor* spp. are still unavailable, except for a few studies that had reviewed its conservation status, conducted a gut metagenomic analysis, as well as sequenced the transcriptome of *T. tambra* (Lau et al., 2021a; Lau et al., 2021b; Lau et al., 2021c). Genomic and transcriptomic sequencing of *Tor* species is necessary to provide a more powerful tool that continues to resolve and address questions of species identification, evolutionary biology, morphological variations, sequences related to sex differentiation, growth, reproduction, and immune, which is helpful for further conservation of *Tor* species (Jaafar et al., 2021).

The rapid growth and expansion of the aquaculture industry had improved fish production as compared to traditional fishing. *T. tambroides* is one of the notable species found in the aquaculture industry due to its high nutritional value and unique flesh taste. However, it has been found that the genetic variability decreases in captivity-held fishes, causing them to be more vulnerable to infections. Thus, it is necessary to comprehend the fish immune system and its underlying mechanisms in response to its exposure to ecotoxicological chemicals (Lim et al., 2018; Lim et al., 2021b). However, in general, the studies of the fish immune response are limited due to its complexity and lack of suitable reagents for classical immunological assays (Salinas & Magádan, 2017). Similarly, as a slow grower fish, it is necessary to venture into the growth-related aspect of the *T. tambroides* genome as well, which is in line with the goal of fish farming of this species which faces knowledge-scarcity on growth improvement to date. Omics approaches, including genomics, transcriptomics, proteomics, and metabolomics, can develop high throughput outcomes and facilitate more novel findings.

In the recent years, with the booming of long-read sequencing technologies (Heather & Chain, 2016), the implementation of combining both Illumina reads and Nanopore/PacBio reads can be found in several studies (Austin et al., 2017; Tan et al., 2018; Lim et al., 2021c; Lim et al., 2022). Integration of short but accurate Illumina reads with long but less accurate Nanopore/PacBio reads could enhance the completeness of the assembled genome than assemblies based on Illumina reads only (Austin et al., 2017; Tan et al., 2018; Lim et al., 2021c; Lim et al., 2022). Thus, in this study, we sequenced and reported on the first genomic data of *T. tambroides*. We also functionally annotated the genome of *T. tambroides* to KEGG, GO, and COG databases and further identified its immune-related genes. Furthermore, we inferred the phylogenetic relationship of *T. tambroides* with other Actinopterygii fishes based on the BUSCO supermatrix data. It is hoped that these data generated from this study can be channeled for the improvement studies that are driven along with the conservation endeavors of this fish species.

## Experimental Design, Materials and Methods

### Sampling and DNA extraction

An adult *T. tambroides* (voucher ID: ASD03018) was sampled from a local aquaculture farm, with its locality reported in previous studies (Lau et al., 2021c; Lim et al., 2021). The fish had been deposited as voucher specimens in the fish museum located at the Faculty of Resource Science and Technology, Universiti Malaysia Sarawak. The fish was euthanised, and 50 mg of its muscle tissue was used for genomic DNA extraction using the DTAB-CTAB DNA extraction kit (GeneReach Biotechnology Corp) according to the manufacturer’s instructions.

### Species Verification

Cytochrome c oxidase subunit I (COI) gene was amplified, and the PCR product was purified and sent for Sanger sequencing. The sequences were analysed using NCBI BLASTn and found to possess 100% similarity with the mitogenome sequences of *T. tambroides* reported by Lim et al. (2021) with the GenBank accession number MW471071.1.

### Library Construction and Whole Genome Sequencing

Approximately one ug of gDNA was sheared to 350 bp using a Bioruptor and directly used for PCR-free library preparation using the NEB Ultra Illumina library preparation kit (NEB, Ipswich, MA). The library was quantified with a Qubit (Invitrogen) and sequenced on a NovaSEQ6000 (Illumina, San Diego, CA) with 2 x 150 bp run configuration. Similarly, for Nanopore sequencing, one ug of unsheared gDNA was used as the input for LSK109 library preparation (Oxford Nanopore, UK) according to the manufacturer’s instructions. The library was sequenced on two MinION flowcells. Nanopore reads were base-called from their fast5 files using Guppy version 4.4.1 (high accuracy mode).

### Sequence Data Processing and Assembly

Illumina reads were trimmed with fastp (Chen et al., 2018), while the Nanopore reads were trimmed with porechop (Wick et al., 2017). The trimmed Nanopore reads and paired-end Illumina reads were used to perform a hybrid *de novo* assembly using Wengan v0.2 (Di Genova et al., 2021). Jellyfish v2.3.0 was used to obtain a frequency distribution of k-mer counting with the clean reads, producing kmer frequency distributions of 31-mers (Marçais & Kingsford, 2011). These histograms were subsequently processed using GenomeScope, which estimates genomic size, repeat content and heterozygosity via kmer-based statistical approach (Vurture et al., 2017). QUAST v5.0.2 was used to evaluate various metrics of the *T. tambroides* genome (Mikheenko et al., 2018) using the parameter: read length 150 bp and max K-mer coverage 1,000. BUSCO v5.2.2 was used to evaluate the completeness of the assembled *T. tambroides* genome based on the single-copy orthologs represented in the actinopterygii_odb10 database (Manni et al., 2021).

### Detection of Repetitive Sequences and Prediction of Protein-Coding Genes

Repetitive sequences in the assembled genomes were identified and masked using RepeatModeler2 and RepeatMasker, respectively. Previously generated transcriptomic reads (Lau et al., 2021b) were aligned to the repeat-masked genome assembly using HiSAT2 (Kim et al., 2015). The transcriptome alignment BAM file and the repeat-masked genome assembly were used as the input for protein-coding gene prediction in BRAKER2 (Bruna et al., 2021).

### SSR Analysis

Simple sequence repeats (SSRs) analysis was identified using Kmer-SSR (https://github.com/ridgelab/Kmer-SSR) (Pickett et al., 2017) on the processed reads of the *T. tambroides* genome. Dinucleotides, trinucleotides, tetranucleotides, pentanucleotides and hexanucleotides were included in the SSRs analysis. Only SSRs with at least four repeats were selected for the study.

### Functional Annotation of Protein-Coding Genes

The predicted protein sequences were functionally annotated to EggNOG mapper (evolutionary genealogy of genes: Non-supervised Orthologous Groups) with a minimum Evalue of 0.001. Functional annotation of genes was performed by mapping against three public databases, GO (Gene ontology), KEGG (Kyoto Encyclopedia of Genes and Genomes), and COG (the Clusters of Orthologous Groups).

In addition, predicted protein sequences were mapped against several previously published growth- and immune-related genes based on literature review (Overturf et al., 2010; Palti, 2011; Vagner & Santigosa, 2011; Zhenzhen et al., 2014; Hu et al., 2016; Ma et al., 2016; Chandhini & Rejish, 2019; Lin et al., 2019; Dam et al., 2020; Damzamnn et al., 2020; Guan & Qiu, 2020). 82 growth- and 31 immune-related genes (113 genes in total) were downloaded from the NCBI GenBank database (https://www.ncbi.nlm.nih.gov/) and used as references. Subsequently, the genes were further filtered based on a stringent E-value cutoff of 10^-10^.

### Ortholog Inference and Phylogenetic Construction

The *Tor tambroides* genome was assembled in this study, while other genome sequences used were obtained from NCBI (National Center for Biotechnology Information) and summarised in Table 1. A total of 50 species genomes together with 2 outgroups (additional species from Coelacanthiformes and Acipenseriformes) were included.

**Table 1.**
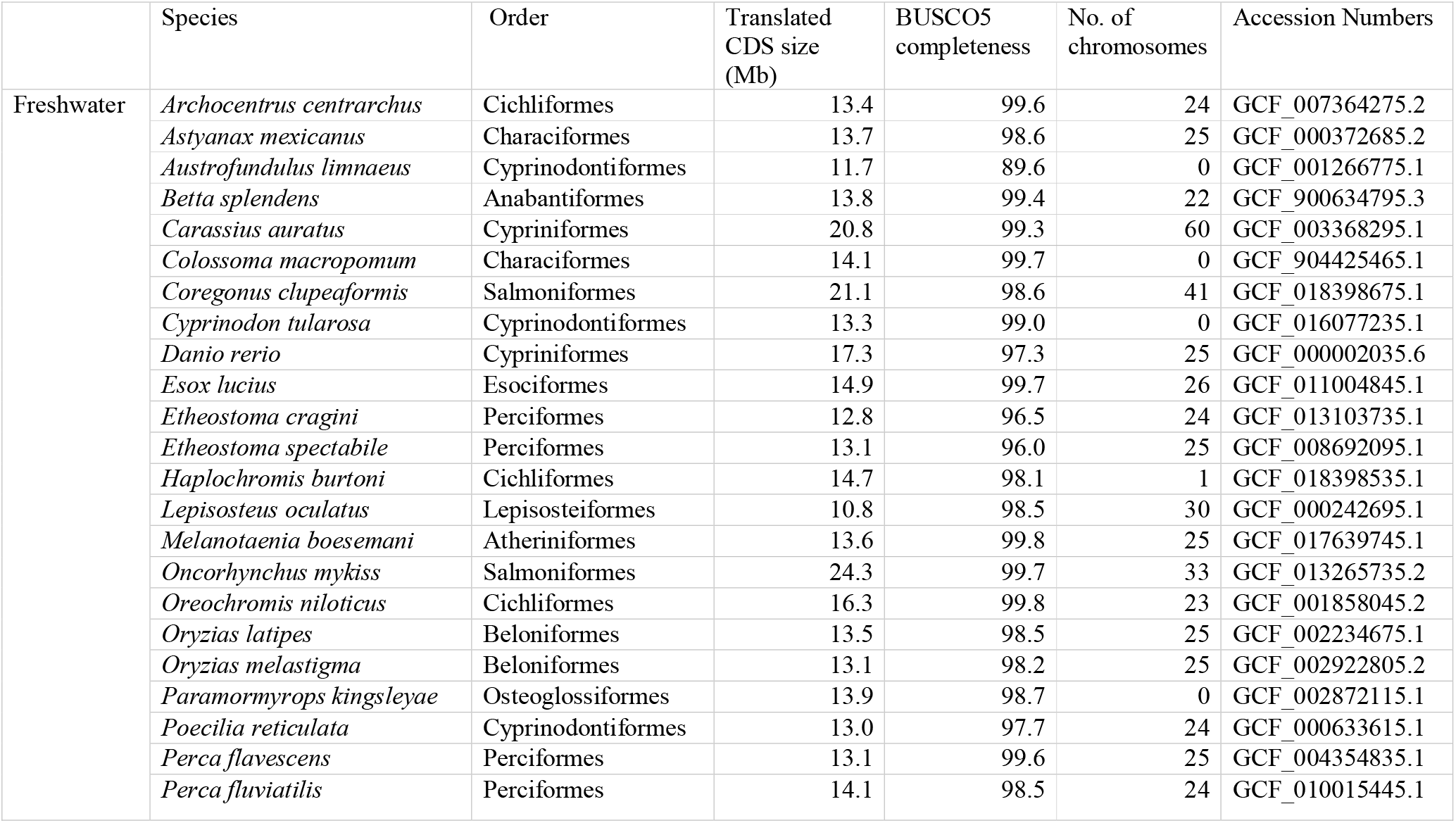
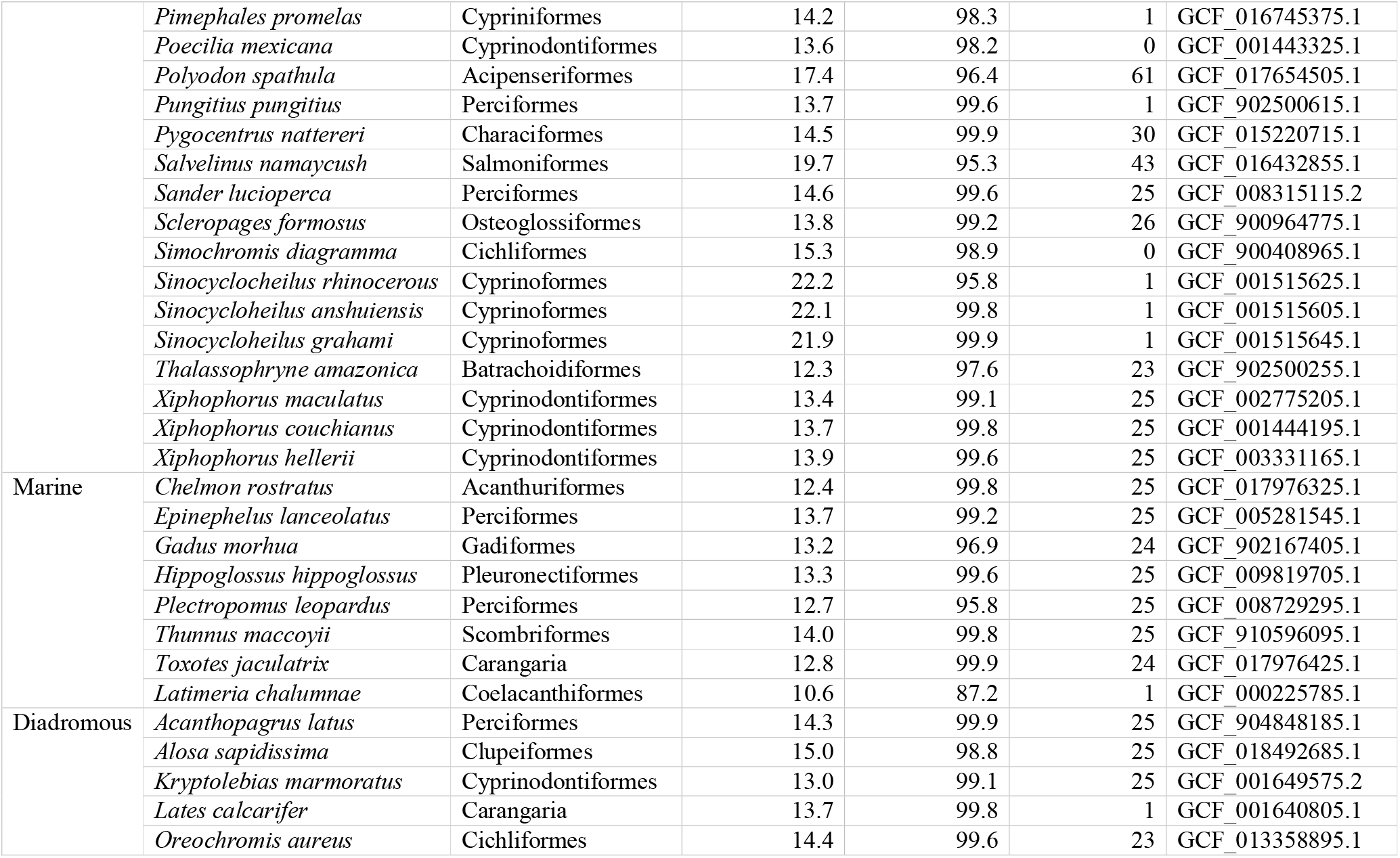
Species found within class Actinopterygii obtained from NCBI were used in this study to build the Maximum-likelihood (ML; IQ-TREE) tree to infer the phylogenetic relationship of these fishes with *T. tambroides*. A total of 50 species with two outgroups were reported and classified into their respective residing environment, including freshwater, marine, or diadromous.

We had identified single-copy orthologous genes in each genome using BUSCO v5.2.2 (Benchmarking Universal Single-Copy Orthologs) (Manni et al., 2021) across all 53 Actinopterygii species. BUSCO was run with orthologs in actinopterygii_odb10 (updated 2021-02-19) using default parameters. All BUSCOs found in a single-copy and multi-copy for each species were used for phylogenetic analysis. Subsequently, BUSCO sequences were individually aligned with MUSCLE (Multiple Sequence Comparison by Log-Expectation) across all 53 species (Edgar, 2004). Gaps and unmatched sites were removed from the resulting multiple sequence alignments (MSAs) using trimAl (Capella-Gutiérrez et al., 2009). These trimmed MSAs were concatenated into a supermatrix. A maximum-likelihood (ML) phylogenetic tree was inferred from the supermatrix using IQ-TREE (Nguyen et al., 2015). Molecular Evolutionary Genetic Analysis (MEGA) v11.0.10 was used to analyse and view the generated phylogenetic tree (Kumar et al., 2018)

The complete genome sequence of *T. tambroides* reported in this study is available under NCBI Bioproject PRJNA708136 (https://submit.ncbi.nlm.nih.gov/subs/bioproject/SUB9225867/overview). The sequencing reads in this study are also available under NCBI SRA entry SUB10492101 (https://submit.ncbi.nlm.nih.gov/subs/sra/SUB10492101).

### Ethics Statement

All experiments comply with ARRIVE guidelines and were carried out in accordance with the U.K. Animals (Scientific Procedures) Act, 1986 and associated guidelines, EU Directive 2010/63/EU for animal experiments, or the National Institute of Health guide for the care and use of Laboratory animals (NIH Publications No. 8023, revised 1978).

## Results & Discussion

### Characterisation of *T. tambroides* and its genome sequencing

The genomic sequencing and assembly statistics of the target *T. tambroides* in this study were summarized in Table 2. The total contig length is 1,235,136,976 bp with 44,726 contigs. The longest contig length recorded is 445,922 bp. The contig GC content documented in this study is 36.55%.

**Table 2.** Genomic sequencing and assembly statistics

|  |  |
| --- | --- |
| Total contig length | 1,235,136,976 bp |
| Number of contigs | 44,726 |
| Longest contig length | 445,922 bp |
| Contig N50 length | 44,262 |
| Contig GC (%) content | 36.55 |
| BUSCO Completeness (Actinopterygii_odb10) |  |
| Actinopterygii_odb10: Complete BUSCOs | 84.3% (3069) |
| Complete and single copy BUSCOs | 39.2% (1427) |
| Complete and duplicated BUSCOs | 45.1% (1642) |
| Missing BUSCOs | 8.3% (301) |

The assembled clean reads of the *T. tambroides* genome were subjected to K-mer analysis using Jellyfish software and visualised using GenomeScope (K-mer:31), as shown in Figure 1. The y-axis had demonstrated the amount of K-mers found at each corresponding depth on the x-axis. There is no heterozygous peak generated at 23, and a low heterozygosity level of 0.194% was recorded. The depth of the homozygous peak can be observed at 46, which accounts for the identical 31-mers from both strands of DNA. The kmer-based statistical approach had revealed that 24.675% of the genomic content is repeated whereas 75.325% of the content is unique. The genome size of *T. tambroides* can be predicted by division of K-mer number over K-mer depth. The k-mer number detected in this study is 69 Gb. Therefore, the expected genome size is predicted as 1.5 Gb.

**Figure 1.**
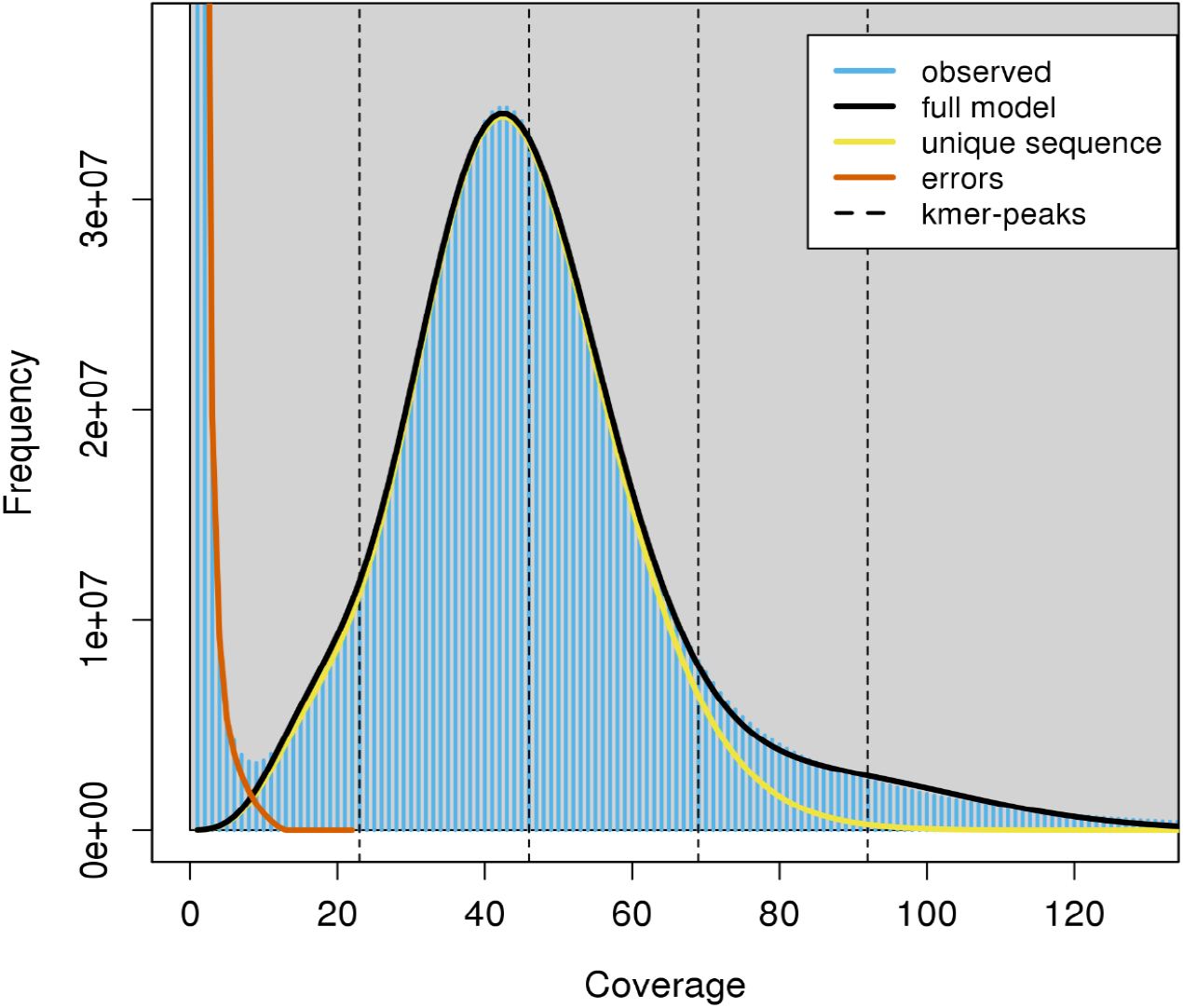
Estimation of genome size, repeat content and heterozygosity by GenomeScope, based on 31-mers (read length = 150 bp; kmer max coverage at 1000).

The *T. tambroides* genome size of 1.5 Gb was found to locate within the range of *Cyprinidae* genome sizes, including grass carp (*Ctenopharyngodon idella*) and common carp (*Cyprinus carpio*) at 1.07 Gb (Wang et al., 2015) and 1.7 Gb (Xu et al., 2014), respectively. Besides, its GC% content reported (36.55%) was slightly lower than the 37.4% and 37.3% seen in *C. idella* and *C. carpio*. In general, a more outstanding GC content was found in seawater (usually above 40%) than freshwater fish (less than 40%), and also in migratory than a non-migratory species (Lu & Luo, 2020). Thus, it is suggested that the genomic GC content may be influenced by different living environments (Lu & Luo, 2020). Besides, the contrary relationship between genomic size (1.5 Gb) and GC content (36.55%) was shown in genome of *T. tambroides*. However, such assertion was insignificant based on a study reviewing 14 species, thus suggesting the collection of more genomic data for further validation (Lu & Luo, 2020).

Repetitive elements accounted for 39.79% in *T. tambroides* genome, which is likely to be associated with its genome duplication. It is found to be in accordance to the repetitive elements found across order Cypriniformes species reporting to be around 35% to 40%, including *Danio rerio, Cyprinus carpio, Sinocycloheilus graham, Sinocycloheilus rhinocerous, Sinocycloheilus anshuiensis* and *Pimephales promelas* (Yuan et al., 2018). Within teleost fish, researchers had found out that the expansion of repetitive elements may be the factor for the expansion of fish genome size as observed across 52 teleost species (Yuan et al., 2018). Having a high repetitive element content within the genome could speed up the generation of novel genes for adaptation purposes. However, an excessive amount of it would bring to abnormal combination and splicing, resulting in unstable genomes (Hong, 1998). In short, it is discouraged for unlimited growth of repetitive elements with its genomic size but limited to certain levels and shaped under specific natural selection (Yuan et al., 2018).

### SSR Analysis

A total of 392,346 SSRs had been successfully identified from the genome of *T. tambroides*. Four types of SSRs had been tabulated with their length, total counts, average length and distribution (Table 3) while Figure 2 is showing the composition of each type of SSR within the genome of *T. tambroides*. The dinucleotide repeats had covered up to 70% of the entire SSR population, encompassing 275,715 SSRs. The trinucleotide and tetranucleotide repeats counts are 59,978 (15%) and 49,089 (13%), and they are ranked second and third out of all the four SSRs. The remaining 2% SSRs were 7,388 pentanucleotide repeats found within the genome of *T. tambroides*. The top three highest frequency dinucleotide repeats SSRs are AT/TA (1,058,829), AC/GT (71,569) and CA/TG (70,249) composing up to 89% of the entire dinucleotide SSRs (Figure 3). The top three trinucleotide repeats covered up to one-third of the trinucleotide SSR repeats, namely AAT/ATT, TTA/TAA and TAT/ATA. As for the tetranucleotide repeats SSRs, the AGAT/ATCT, GATA/TATC and TCTA/TCGA counts were 6,209, 5,386 and 3,826 respectively, covering up to 30% of the entire tetranucleotide SSRs. Correspondingly, 18% of the pentanucleotide repeat SSRs were AAAAT/ATTTT (6.42%), ATAAT/ATTAT (6.13%) and TTTTA/TAAAA (5.96%) respectively. Overall, the dinucleotides SSRs, namely AT/TA (26.97%), AC/GT (25.96%) and CA/TG (17.90%) had occupied more than half of the SSR population found within the genome of *T. tambroides*.

**Table 3.**
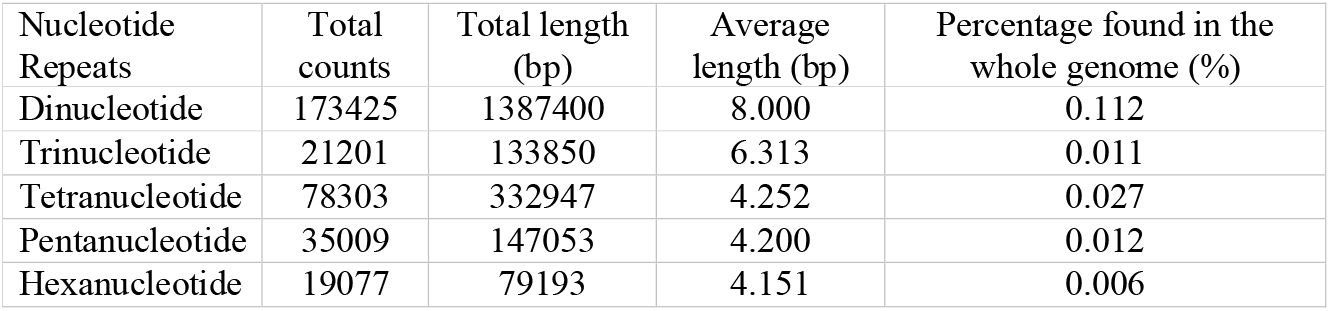
The number, total length and average length of five different types of SSRs found within the genome of *T. tambroides*.

| Nucleotide Repeats | Total counts | Total length (bp) | Average length (bp) | Percentage found in the whole genome (%) |
| --- | --- | --- | --- | --- |
| Dinucleotide | 173425 | 1387400 | 8.000 | 0.112 |
| Trinucleotide | 21201 | 133850 | 6.313 | 0.011 |
| Tetranucleotide | 78303 | 332947 | 4.252 | 0.027 |
| Pentanucleotide | 35009 | 147053 | 4.200 | 0.012 |
| Hexanucleotide | 19077 | 79193 | 4.151 | 0.006 |

**Figure 2.**
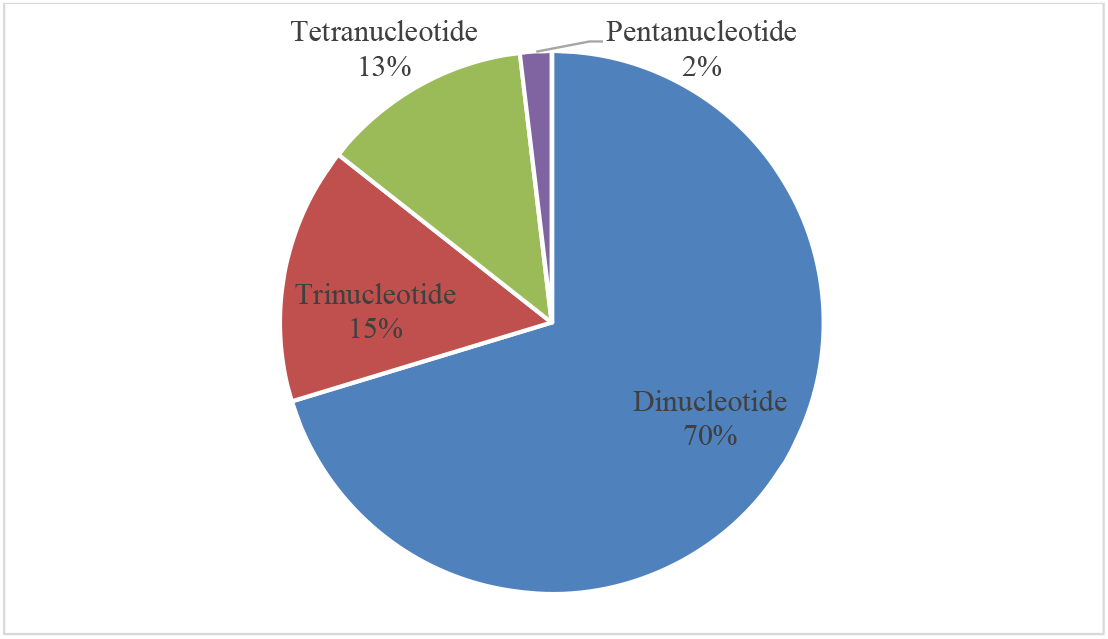
The composition of dinucleotide, trinucleotide, tetranucleotide, pentanucleotide and hexanucleotide SSRs identified from the genome of *T. tambroides*.

**Figure 3.**
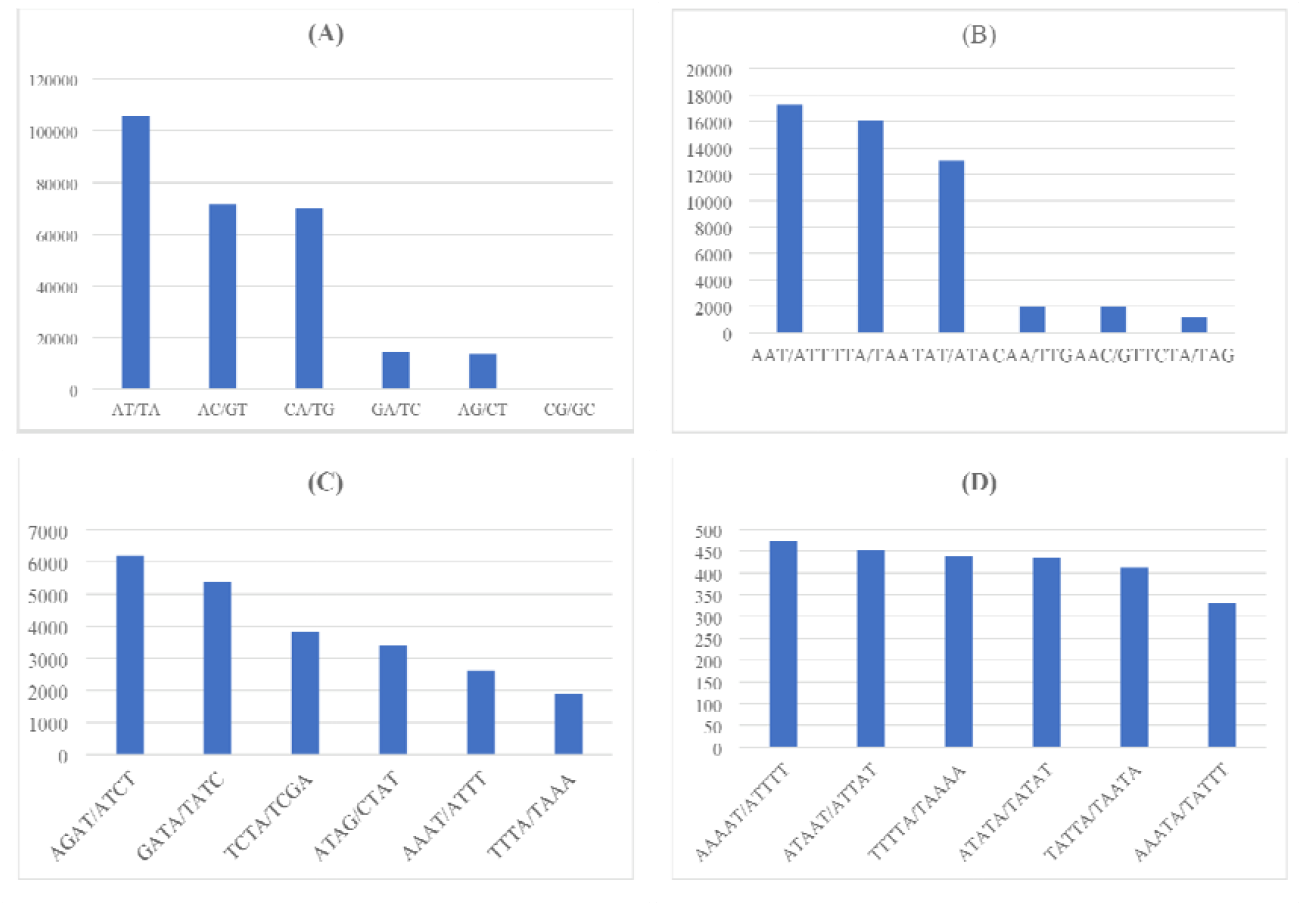
SSRs percentage graph with selected top six frequency SSRs from each group: (A) dinucleotide repeats, (B) trinucleotide repeats, (C) tetranucleotide repeats, (D) pentatetranucleotide repeats.

SSRs allow faster adaptation to environmental stress through increment of DNA quantity and raw material for adaptative evolution during genome evolution. Thus, it can be said the mutation rate of microsatellite is dependent on the repeated unit length with a more common observation of mono- and dinucleotide repeats as compared to other repeats with longer lengths due to respective stability (Schlotterer, 2000). The frequency of tetranucleotide was higher than trinucleotide within the genome of *T. tambroides*, which was shown in other ray-finned fishes as well (Lei et al., 2021). Less occurrence of trinucleotide SSRs repeats can be due to its attribution as a triplet code to form part of the gene and also the presence of a mismatch repair system in the exonic region to maintain greater trinucleotide repeats (Lei et al., 2021).

SSR repeats with poly (A/T) tracts were found in a greater abundance than repeats with poly (G/C) tracts in other ray-finned fishes across all types of SSRs including dinucleotide, trinucleotide, tetranucleotide and pentanucleotide (Lei et al., 2021). The higher frequency of poly (A) can be due to the re-integration of the processed genes from mRNA back into the genome with an attached poly (A) tail, while poly (G/C) is not included in this integrative mechanism (Lei et al., 2021). In addition, greater poly (A) occurrence can be explained through the formation of pseudogenes and its necessity in the universal retrotransposon (Toth et al., 2000; Borodulina et al., 2016). The (GC) repeats are more stable than (AT) repeats thus increasing the difficulty to be slipped during replication (Gur-Arie et al., 2000).

Dinucleotides AT/TA is the common microsatellites repeat found across the fish genomes (Lei et al., 2021) which is observed within the genome of *T. tambroides* as well. Furthermore, for trinucleotide repeats, the occurrence of (CCG) n (16 counts) and (ACG) n (2572 counts; 5%) repeats were rare in *T. tambroides* as well. This phenomenon can be explained by the presence of the highly mutable CpG dinucleotide within the motif due to methylation (Lei et al., 2021). While in tetranucleotide repeats, the G+C content of SSRs was observed in a lower frequency because of its influence on the mutation rate as there is no statistical significance between 25% G+C content but each was significant difference from the 50% G+C repeat content (Eckert et al., 2002).

The repetitive element found within the genome of *T. tambroides* was reported as 4.15%. It had been reported previously that the expansion of repetitive elements would cause a further expansion in the fish genome Therefore, it can be said that the fish genome size is positively correlated with the repetitive elements (Yuan et al., 2018). However, a study reviewing the SSR across 14 fish species contradicts the statement (Lei et al., 2021). The generation of novel genes for adaptation can be accelerated with the presence of high repetitive element content, for instance in salmon, which is likely to be associated with genome duplication (Lien et al., 2016). Besides, the variation observed within microsatellites can be due to differential selective constraints causing accumulated preference for different microsatellite types (Lei et al., 2021). However, overburden could cause abnormal recombination and splicing, resulting in genome instability (Yuan et al., 2018). In short, it can be concluded that the repetitive elements must be limited to shape under specific natural selection by the environment. Its unambiguous role in genomic function still remains to be explored.

### Functional Annotation of *T. tambroides*

For functional annotation of *T. tambroides* genome, coding region was extracted using Interproscan (Jones et al., 2014), generating 96,736 predicted non-redundant protein sequences. Subsequently, the sequences were annotated using eggNOG mapper to map the predicted protein sequences to GO, KEGG, and COG databases. The sequence length of each unigene ranged from < 100 bp to > 2000 bp (Figure 4), which showed a decreasing trend as the length increases. Table 4 shows the number of unigenes annotated to either GO, KEGG or COG, unigenes annotated to at least one of the databases and all the databases. A total of 42,694 (75.19%), 25,560 (45.01%), and 55,981 (98.58%) of unigenes were annotated to GO, KEGG, and COG databases respectively. Out of a total of 56,785 annotated unigenes, 56,094 (98.78%) of the unigenes had been found to have a significant match to at least one of the databases while 21,878 (38.53%) unigenes portrayed a notable match to all the three databases. Figure 5 illustrates the distribution of unigenes across GO, KEGG, and COG databases.

**Figure 4.**
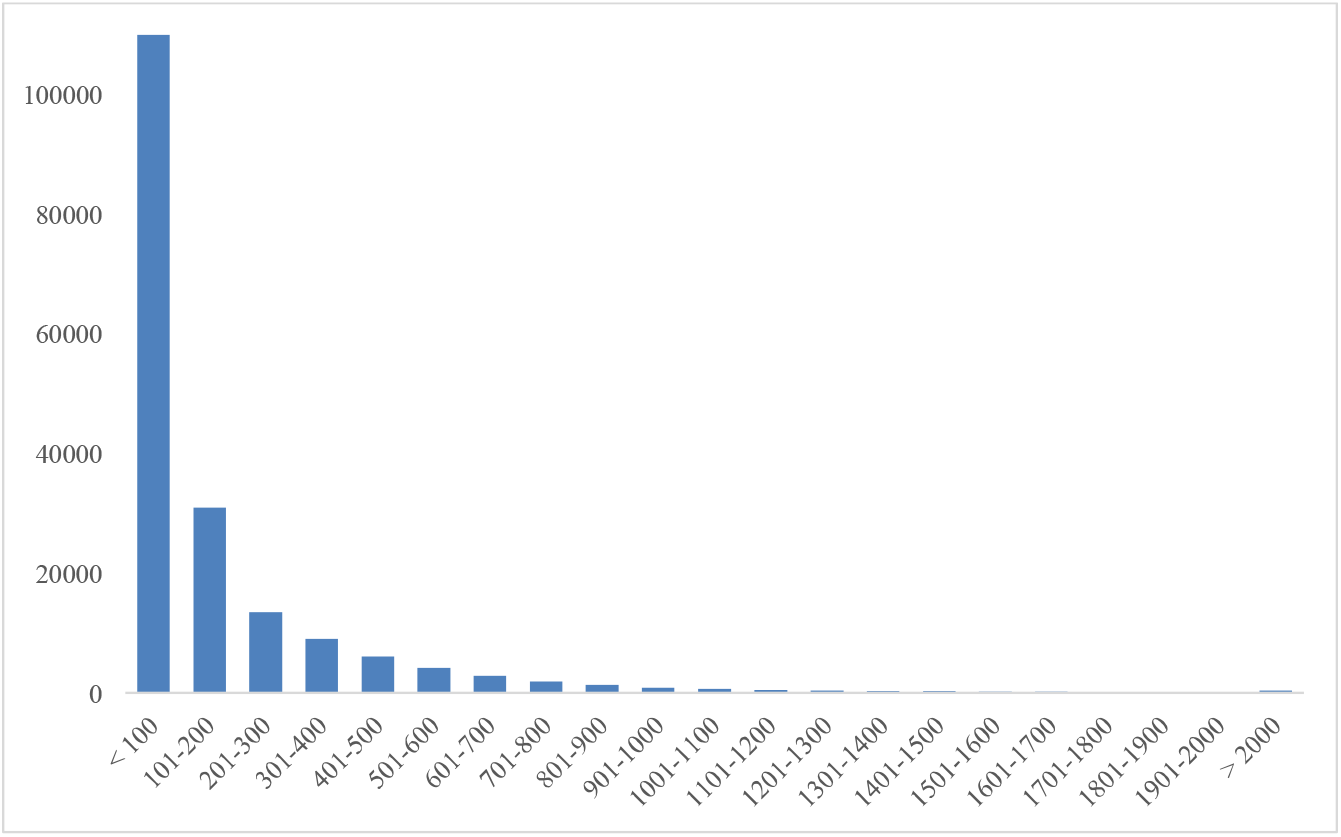
Length distribution of unigenes of *T. tambroides*.

**Figure 5.**
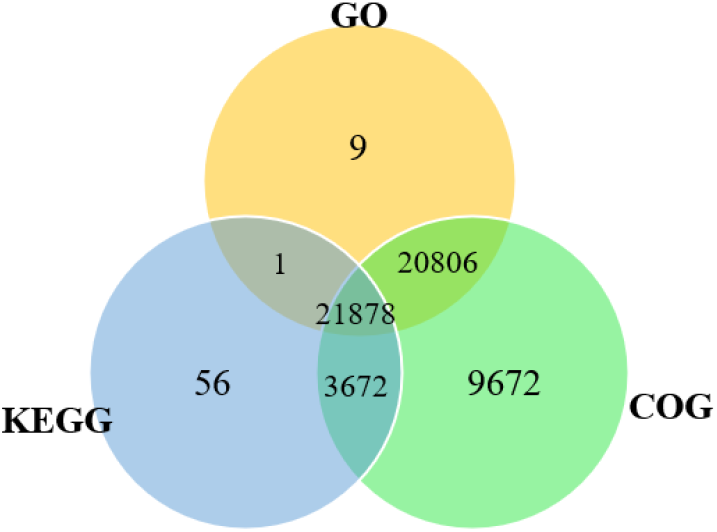
Venn diagram showing differences and similarity of unigenes of *T. tambroides* annotated to GO, KEGG, and COG databases.

**Table 4.** Functional annotation of unigenes to the various database.

| Database | Number of Unigenes | Percentage (%) |
| --- | --- | --- |
| GO | 42,694 | 75.19 |
| KEGG | 25,560 | 45.01 |
| COG | 55,981 | 98.58 |
| Annotated in at least one database | 56,094 | 98.78 |
| Annotated in all database | 21,878 | 38.53 |
| All unigenes | 56,785 | 100.00 |

Annotation of *T. tambroides* genome to each main ontology of GO database was shown in Figure 6, including biological process, molecular functions, and cellular components. Under biological process, metabolism (4475; 6.56%) had the greatest count, followed by development (3337; 4.89%) and catalytic activity (2149; 3.15%). On the other hand, a total of 1239 counts (1.82%) were responsible for binding, 761 (1.12%) accounted for transferase activity and 688 (1.01%) accounted for protein binding, under the molecular function category. Furthermore, under the cellular components category, 2139 (3.14%) were accounted for cell organization and biogenesis while 1672 (2.45%) and 1284 (1.88%) were categorized as cell and intracellular respectively.

**Figure 6.**
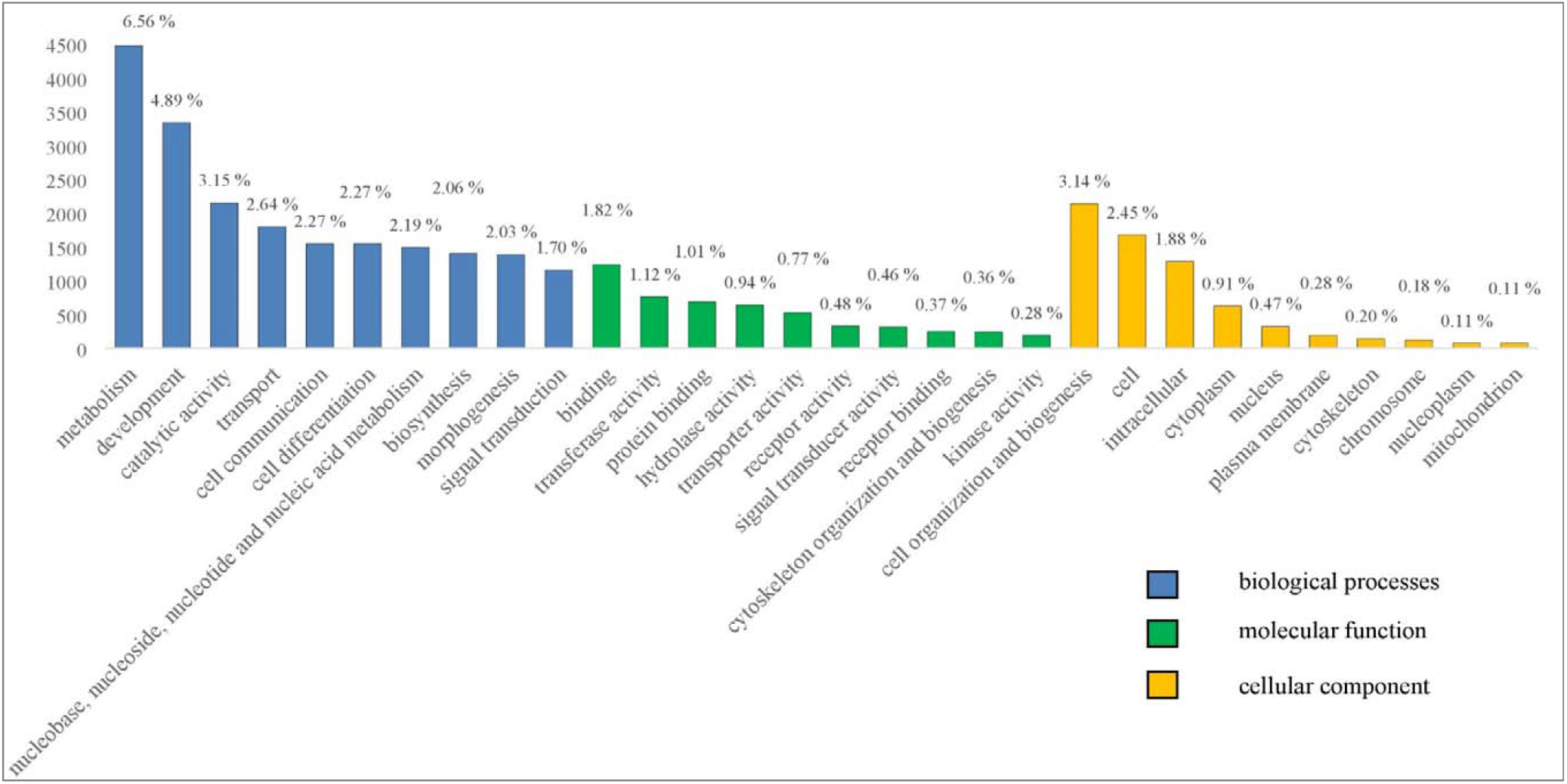
GO classifications

Another annotation was performed towards a widely-used reference database KEGG equipped with multiple pathways for better integration and interpretation of large-scale datasets. *T. tambroides* genome had successfully mapped towards 304 known KEGG pathways (Figure 7), including organismal system, cellular processes, environmental information processing, genetic information processing and metabolism. Out of the five main aforementioned categories, the largest count (36340; 39.4%) is from organismal system whilst genetic information processing (4324; 4.69%) had the lowest count. The categories reported on the greatest number of counts were signal transduction from environmental information processing (19577; 21.22%), endocrine system from organismal system (9415; 8.79%) and immune system (8110; 10.21%) from organismal system. Figure 8 depicts the top ten KEGG cluster components found in each main aforementioned category. The top three largest count can be observed in metabolic pathway (4302; 4.66%), PI3K-Akt signaling pathway (1500; 1.63%) and MAPK signaling pathway (1464; 1.59%) from signal transduction.

**Figure 7.**
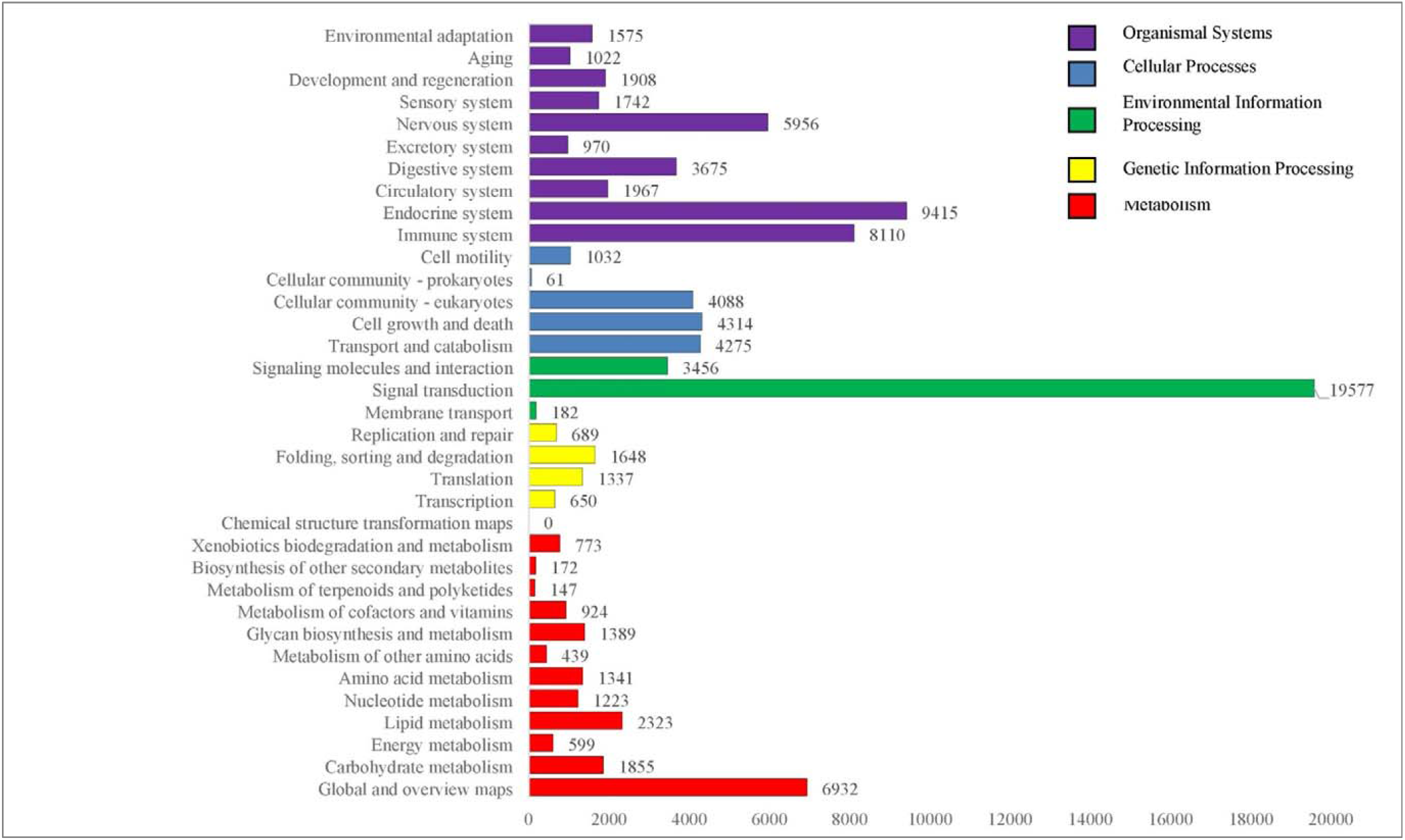
KEGG classifications

**Figure 8.**
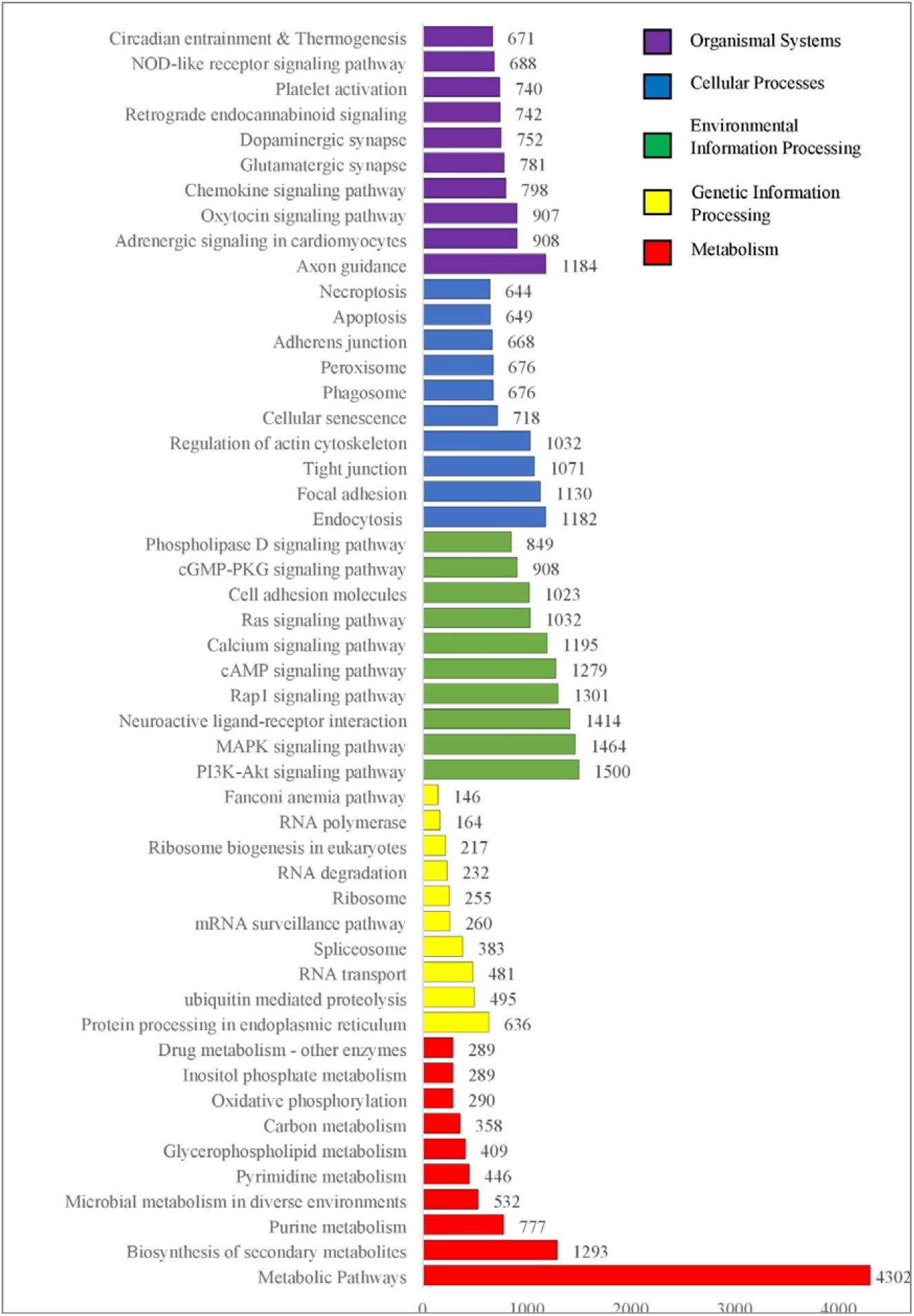
Top 10 KEGG classification

Figure 9 illustrates the classification of 96,736 predicted protein sequences towards COG database consisting of clusters of orthologous groups. There were altogether 25 COG classifications and can be grouped under four main clusters: information storage and processing (8906; 15.17%), cellular processes and signaling (25248; 43.01%), metabolism (8402; 14.31%) and poorly characterized (16152; 27.51%). Among all the clusters, greater counts were originated from function unknown (16152; 27.51%), followed by signal transduction mechanism (14284; 24.33%) and transcription (4455; 7.59%). Overall, the genome of *T. tambroides* is enriched with gene families in the categories of signal transduction, endocrine system, immune system and metabolic pathways. It is found to be consistent with the common carp genome (Xu et al., 2014) and Javan mahseer transcriptome (Lau et al., 2021b).

**Figure 9.**
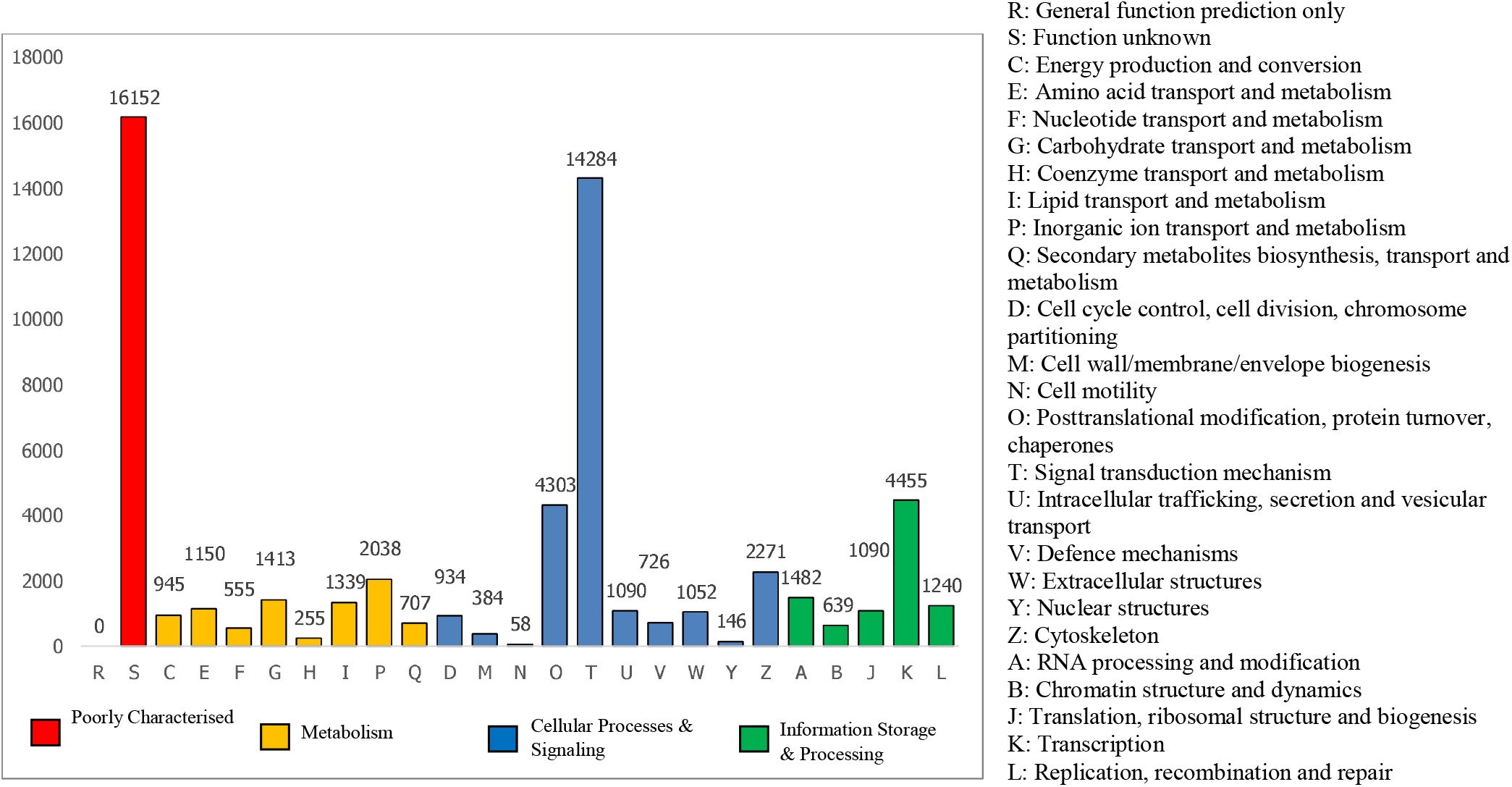
COG classifications

### Immunity-related Annotation

To characterize a comprehensive defense landscape of *T. tambroides* against pathogenic infections, the genomic sequences of *T. tambroides* had been functionally annotated to identify pathways and genes associated with the fish immunity. GO classification had revealed on 14,489 unigenes (63.15%) were functionally annotated to immune-related GO terms. Among the annotated categories, metabolism (4475; 30.89%), stress response (1824; 12.59%) and protein metabolism (1135; 7.83%) had reported on the greatest counts. KEGG pathway analysis had revealed 41 immune-related pathways, including MAPK signaling pathway, Toll-like receptor (TLR) signaling pathway, Wnt signaling pathway, NOD-like receptor signaling pathway and so on (Figure 10) (Zhenzhen et al., 2014).

**Figure 10.**
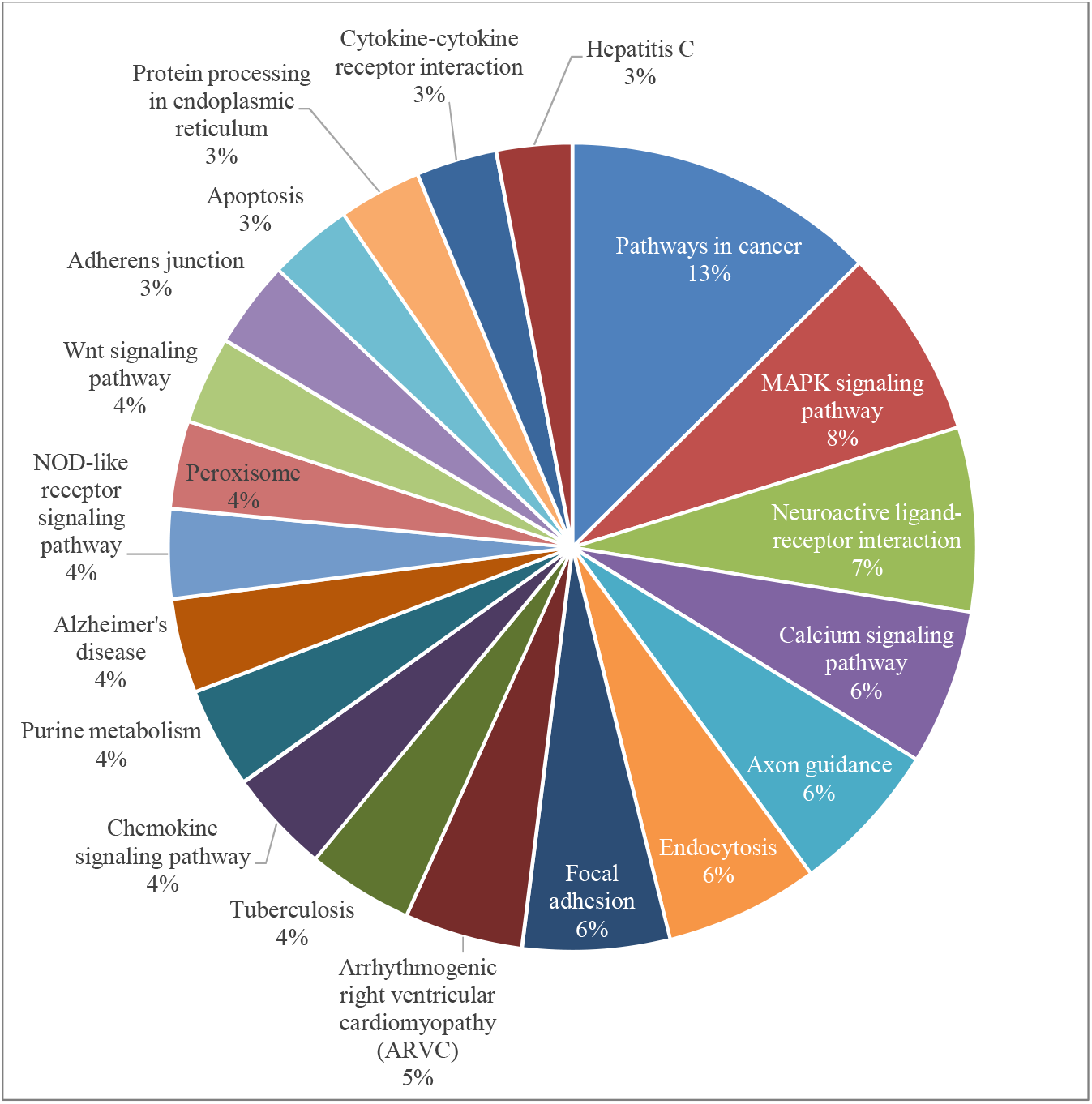
Top 20 annotated KEGG immune pathway of *T. tambroides* genome.

Among all the components in the fish immune system, TLR family is an essential type of pattern-recognition receptors expressing on antigen-presenting cells, involved in innate immune response and the subsequent promotion of adaptive immunity (Akira et al., 2006). We had successfully identified 11 different *TLRs* and 12 *TLR* genes matched to the genome of *T. tambroides* (Table 5). The major histocompatibility complex (MHC) are the important molecules for the recognition of foreign substances via binding peptide fragments from pathogens and presenting them for T cell elimination (Neefjes et al., 2011). Both MHC class I and II had been identified in the genomic dataset as well. MHC genes serve as a candidate disease resistance marker due to their highly polymorphic characteristics in teleost (Langefors et al., 2001). Both TLR and MHC were also found within the transcriptomic dataset of *Trachinotus ovatus* (Zhenzhen et al., 2014). Thus, it is believed that further analysis of both genes will be able to provide insights into the immune system of *T. tambroides* as well as other teleosts.

**Table 5.**
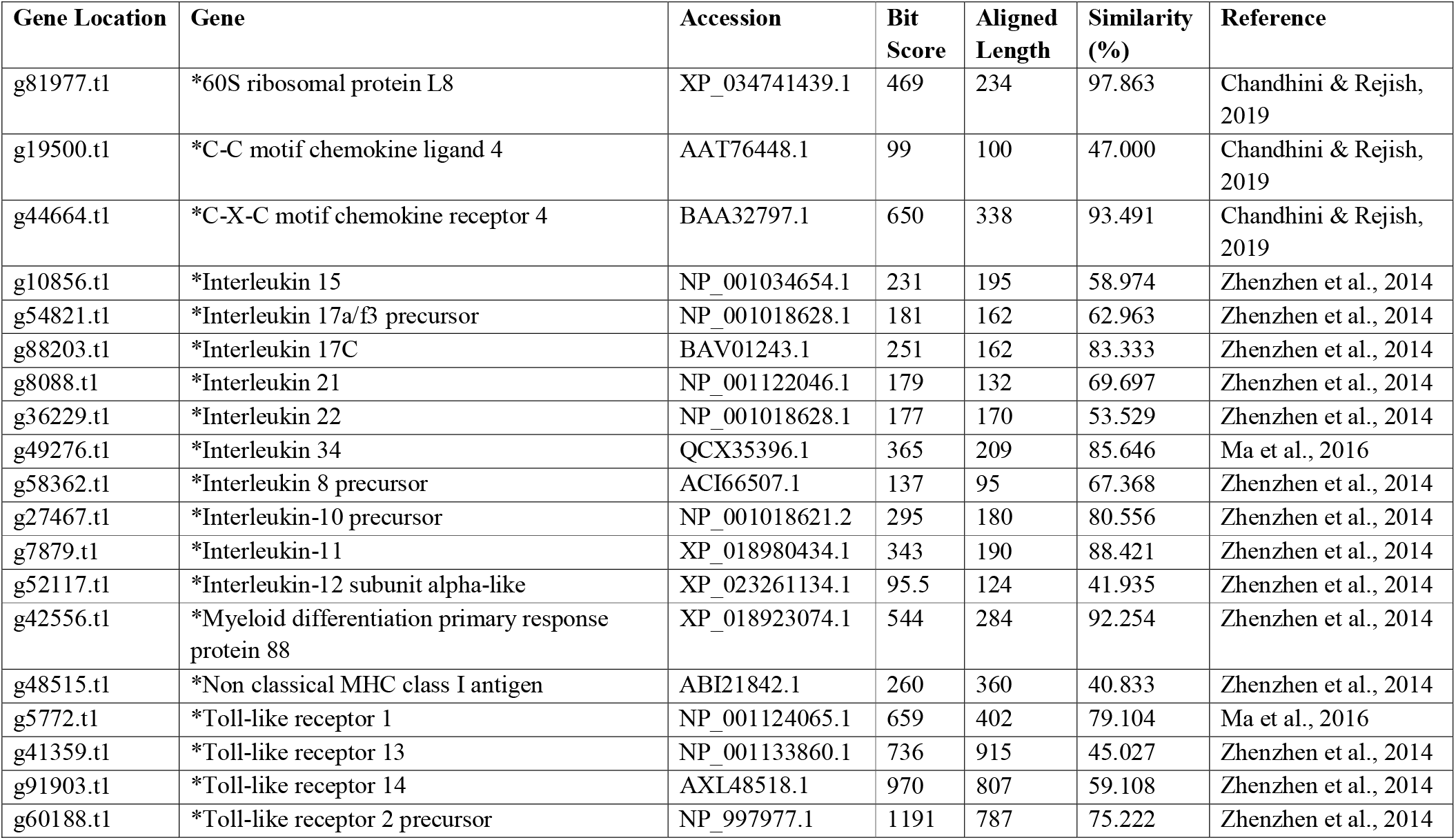
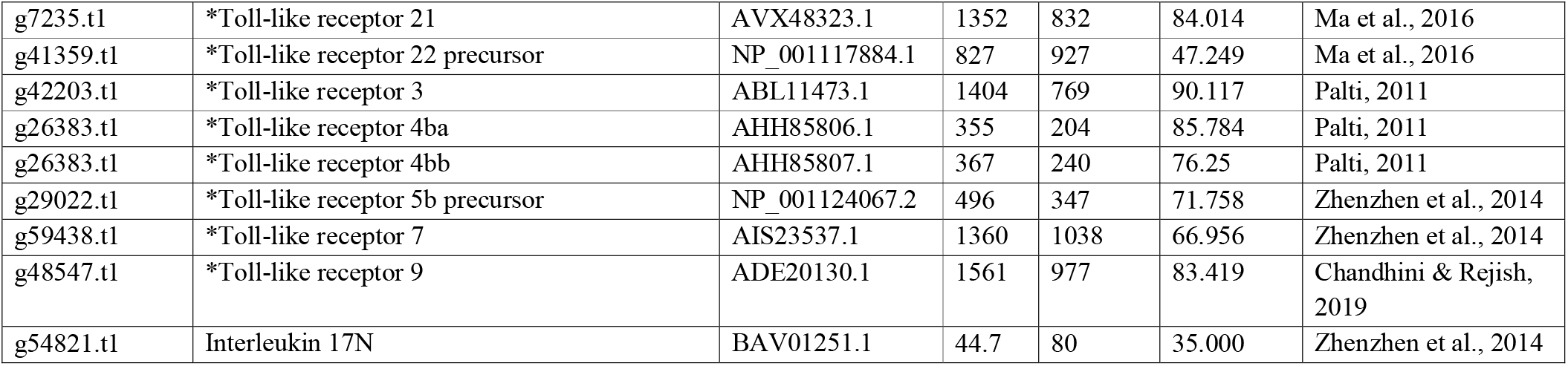
Immune-related genes. Genes marked with * asterisk sign were genes selected after e-value cutoff while best parameters were inputted for genes that did not pass the cutoff filter.

| Gene Location | Gene | Accession | Bit Score | Aligned Length | Similarity (%) | Reference |
| --- | --- | --- | --- | --- | --- | --- |
| g81977.t1 | *60S ribosomal protein L8 | XP_034741439.1 | 469 | 234 | 97.863 | Chandhini & Rejish, 2019 |
| g19500.t1 | *C-C motif chemokine ligand 4 | AAT76448.1 | 99 | 100 | 47.000 | Chandhini & Rejish, 2019 |
| g44664.t1 | *C-X-C motif chemokine receptor 4 | BAA32797.1 | 650 | 338 | 93.491 | Chandhini & Rejish, 2019 |
| g10856.t1 | *Interleukin 15 | NP_001034654.1 | 231 | 195 | 58.974 | Zhenzhen et al., 2014 |
| g54821.t1 | *Interleukin 17a/f3 precursor | NP_001018628.1 | 181 | 162 | 62.963 | Zhenzhen et al., 2014 |
| g88203.t1 | *Interleukin 17C | BAV01243.1 | 251 | 162 | 83.333 | Zhenzhen et al., 2014 |
| g8088.t1 | *Interleukin 21 | NP_001122046.1 | 179 | 132 | 69.697 | Zhenzhen et al., 2014 |
| g36229.t1 | *Interleukin 22 | NP_001018628.1 | 177 | 170 | 53.529 | Zhenzhen et al., 2014 |
| g49276.t1 | *Interleukin 34 | QCX35396.1 | 365 | 209 | 85.646 | Ma et al., 2016 |
| g58362.t1 | *Interleukin 8 precursor | ACI66507.1 | 137 | 95 | 67.368 | Zhenzhen et al., 2014 |
| g27467.t1 | *Interleukin-10 precursor | NP_001018621.2 | 295 | 180 | 80.556 | Zhenzhen et al., 2014 |
| g7879.t1 | *Interleukin-11 | XP_018980434.1 | 343 | 190 | 88.421 | Zhenzhen et al., 2014 |
| g52117.t1 | *Interleukin-12 subunit alpha-like | XP_023261134.1 | 95.5 | 124 | 41.935 | Zhenzhen et al., 2014 |
| g42556.t1 | *Myeloid differentiation primary response protein 88 | XP_018923074.1 | 544 | 284 | 92.254 | Zhenzhen et al., 2014 |
| g48515.t1 | *Non classical MHC class I antigen | ABI21842.1 | 260 | 360 | 40.833 | Zhenzhen et al., 2014 |
| g5772.t1 | *Toll-like receptor 1 | NP_001124065.1 | 659 | 402 | 79.104 | Ma et al., 2016 |
| g41359.t1 | *Toll-like receptor 13 | NP_001133860.1 | 736 | 915 | 45.027 | Zhenzhen et al., 2014 |
| g91903.t1 | *Toll-like receptor 14 | AXL48518.1 | 970 | 807 | 59.108 | Zhenzhen et al., 2014 |
| g60188.t1 | *Toll-like receptor 2 precursor | NP_997977.1 | 1191 | 787 | 75.222 | Zhenzhen et al., 2014 |
| g7235.t1 | *Toll-like receptor 21 | AVX48323.1 | 1352 | 832 | 84.014 | Ma et al., 2016 |
| g41359.t1 | *Toll-like receptor 22 precursor | NP_001117884.1 | 827 | 927 | 47.249 | Ma et al., 2016 |
| g42203.t1 | *Toll-like receptor 3 | ABL11473.1 | 1404 | 769 | 90.117 | Palti, 2011 |
| g26383.t1 | *Toll-like receptor 4ba | AHH85806.1 | 355 | 204 | 85.784 | Palti, 2011 |
| g26383.t1 | *Toll-like receptor 4bb | AHH85807.1 | 367 | 240 | 76.25 | Palti, 2011 |
| g29022.t1 | *Toll-like receptor 5b precursor | NP_001124067.2 | 496 | 347 | 71.758 | Zhenzhen et al., 2014 |
| g59438.t1 | *Toll-like receptor 7 | AIS23537.1 | 1360 | 1038 | 66.956 | Zhenzhen et al., 2014 |
| g48547.t1 | *Toll-like receptor 9 | ADE20130.1 | 1561 | 977 | 83.419 | Chandhini & Rejish, 2019 |
| g54821.t1 | Interleukin 17N | BAV01251.1 | 44.7 | 80 | 35.000 | Zhenzhen et al., 2014 |

### Growth-related Annotation

To tap into the growth-related aspect of the *T. tambroides* genomic landscape, we functionally annotated the genome based on previously characterized growth-related genes. Table 6 shows the mapped BLAST results of immune-related genes on the genome of *T. tambroides* where there were 74 genes labelled with asterisk signs that had passed the Evalue cutoff filter of 10^-10^. KEGG pathway analysis revealed 19 pathways associated with growth, including pathways in cancer, insulin signaling pathway, endocytosis, focal adhesion, and mTOR signaling pathway (Zhenzhen et al., 2014) (Figure 11).

**Table 6.** Growth-related genes. Genes marked with * asterisk sign were proteins selected after e-value cutoff while best parameters were input for the gene that did not pass the cutoff filter.

| Gene Location | Gene | Accession | Bit Score | Aligned Length | Similarity (%) | Reference |
| --- | --- | --- | --- | --- | --- | --- |
| g79685.t1 | *Acetyl-CoA carboxylase alpha (Haplotype 1) | ADT82650.1 | 4772 | 2390 | 95.314 | Guan & Qiu, 2020 |
| g79685.t1 | *Acetyl-CoA carboxylase alpha (Haplotype 2) | ADX43925.1 | 4772 | 2390 | 95.816 | Guan & Qiu, 2020 |
| g44363.t1 | *Acetyl-CoA carboxylase 2 | XP_018962132.1 | 3695 | 1845 | 96.585 | Guan & Qiu, 2020 |
| g54099.t1 | *Acetyl-CoA Acetyltransferase 2 | AAD34966.1 | 751 | 395 | 92.152 | Hu et al., 2016 |
| g1109.t1 | *Alpha actinin 3 | XP_031588180.1 | 1574 | 897 | 82.051 | Damzamn et al., 2020 |
| g59012.t1 | *Adiponectin | NP_001373470.1 | 1574 | 897 | 82.051 | Hu et al., 2016 |
| g16898.t1 | *Alpha-1-Microglobulin/Bikunin Precursor | XP_005451429.1 | 189 | 189 | 44.444 | Dam et al., 2020 |
| g82874.t1 | *Apolipoprotein B | NP_001013332.1 | 407 | 259 | 78.764 | Dam et al., 2020 |
| g83831.t1 | *ATP synthase | BAA82837.1 | 961 | 518 | 93.822 | Dam et al., 2020 |
| g40123.t1 | *ATPase H<sup>+</sup> transporting V1 subunit E1 | AAM34666.1 | 440 | 226 | 96.460 | Guan & Qiu, 2020 |
| g8813.t1 | *G2/mitotic-specific cyclin-B1 | NP_571588.1 | 112 | 215 | 91.071 | Guan & Qiu, 2020 |
| g31308.t1 | *Cell division cycle protein 20 | NP_998245.2 | 919 | 496 | 90.726 | Guan & Qiu, 2020 |
| g3538.t1 | *Conserved Edge Expressed | ABK35126.2 | 548 | 326 | 90.184 | Overturf et al., 2010 |
| g81872.t1 | *Chymotrypsin-like elastase family member 1 | XP_018937349.1 | 461 | 264 | 93.561 | Dam et al., 2020 |
| g49416.t1 | *Creatine kinase M-type | XP_028660838.1 | 616 | 366 | 80.874 | Damzamn et al., 2020 |
| g66522.t1 | *Peroxisomal carnitine O-octanoyltransferase | XP_018953213.1 | 1228 | 612 | 94.935 | Lin et al., 2019 |
| g50598.t1 | *Cathepsin K precursor | NP_001017778.1 | 634 | 333 | 89.189 | Dam et al., 2020 |
| g2135.t1 | *Cathepsin L | AAI08032.1 | 568 | 336 | 76.786 | Dam et al., 2020 |
| g46341.t1 | *25-hydroxycholesterol_7-alpha-hydroxylase | XP_699028.2 | 878 | 506 | 82.016 | Lin et al., 2019 |
| g16441.t1 | *Delta-6_desaturase | AZL94116.1 | 926 | 444 | 100.00 | Vagner & Santiagosa, 2011 |
| g39490.t1 | *Elongation of very long chain fatty acids protein 6 | XP_018957993.1 | 438 | 266 | 82.707 | Dam et al., 2020 |
| g4692.t1 | *Fatty acid-binding protein 10-A | NP_694492.1_ | 240 | 126 | 90.476 | Hu et al., 2016 |
| g23845.t1 | *Fatty acid synthase | ARO92273.1 | 4788 | 2486 | 94.328 | Lin et al., 2019 |
| g76435.t1 | *Farnesyl pyrophosphate synthase isoform X1 | XP_005472704.1 | 608 | 359 | 81.058 | Lin et al., 2019 |
| g19569.t1 | *Forkhead box protein K1 isoform X1 | XP_025764132.1 | 790 | 695 | 64.173 | Lin et al., 2019 |
| g33446.t1 | *G0/G1 switch protein 2 | AVM4087.1 | 215 | 112 | 91.071 | Damzamn et al., 2020 |
| g51734.t1 | *Glucose-6-phosphatase | AVP32214.1 | 649 | 351 | 95.157 | Dam et al., 2020 |
| g87856.t1 | *Growth hormone | CAA43007.1 | 385 | 210 | 95.238 | Ma et al., 2016 |
| g40036.t1 | *Growth hormone receptor | ADZ13484.1 | 490 | 616 | 46.753 | Ma et al., 2016 |
| g40036.t1 | *Growth hormone receptor 1 | AAY86769.1 | 490 | 616 | 46.753 | Hu et al., 2016 |
| g80039.t1 | *Glutathione synthetase | XP_018970229.1 | 883 | 475 | 90.105 | Lin et al., 2019 |
| g71468.t1 | *70-kDa heat shock proteins | AAF70445.1 | 1125 | 634 | 86.278 | Hu et al., 2016 |
| g74743.t1 | *Insulin-like growth factor-binding protein 1 | ACV72066.1 | 524 | 262 | 96.947 | Guan & Qiu, 2020 |
| g58943.t1 | *Insulin-like growth factor-binding protein 2 | ACM47497.1 | 548 | 274 | 97.080 | Guan & Qiu, 2020 |
| g74742.t1 | *Insulin-like growth factor-binding protein 3 | ACM47527.1 | 313 | 173 | 88.439 | Guan & Qiu, 2020 |
| g38084.t1 | *Inositol Monophosphatase 1 | XP_018975133.1 | 571 | 282 | 97.518 | Lin et al., 2019 |
| g85602.t1 | *Insulin-induced_gene_1_protein | NP_956163.1 | 481 | 242 | 97.107 | Hu et al., 2016 |
| g59760.t1 | *Leptin b1 | QEN91938.1 | 337 | 168 | 95.238 | Zhenzhen et al., 2014 |
| g72992.t1 | *Leptin A2 | AGK24956.1 | 309 | 171 | 88.889 | Zhenzhen et al., 2014 |
| g83000.t1 | *Tyrosine-protein kinase JAK2-like | XP_022620925.1 | 1602 | 967 | 78.904 | Zhenzhen et al., 2014 |
| g61842.t1 | *Lipopolysaccharide binding protein | NP_001118057.1 | 672 | 469 | 69.936 | Hu et al., 2016 |
| g55290.t1 | *Hepatic triacylglycerol lipase | XP_018956861.1 | 964 | 498 | 92.169 | Dam et al., 2020 |
| g49598.t1 | *Myocyte Enhancer Factor 2 | BAA33567.1 | 372 | 475 | 51.158 | Overturf et al., 2010 |
| g21088.t1 | *Myostatin 1 | AJF48833.1 | 515 | 351 | 68.661 | Overturf et al., 2010 |
| g21088.t1 | *Myostatin 2 | AJF48834.1 | 726 | 366 | 94.536 | Overturf et al., 2010 |
| g94911.t1 | *Myogenic factor 5 | AWG41988.1 | 486 | 240 | 97.083 | Ma et al., 2016 |
| g28564.t1 | *Neuropeptide Y receptor Y7 | NP_001007219.1 | 650 | 372 | 88.710 | Zhenzhen et al., 2014 |
| g34597.t1 | *Cytochrome P450 | AAK37960.1 | 711 | 505 | 66.535 | Hu et al., 2016 |
| g21266.t1 | *Paired box protein 7 isoform X1 | XP_013988550.1 | 580 | 496 | 62.903 | Overturf et al., 2010 |
| g37657.t1 | *Cytosolic phospholipase A2 gamma-like | XP_018952444.1 | 660 | 558 | 60.394 | Dam et al., 2020 |
| g45170.t1 | *Proopiomelanocortin | AAM93491.2 | 387 | 222 | 83.333 | Zhenzhen et al., 2014 |
| g68580.t1 | *Peroxisome proliferator-activated receptor alpha | CAJ76702.1 | 698 | 468 | 73.077 | Hu et al., 2016 |
| g41063.t1 | *Peroxisome proliferator-activated receptor beta | ACR15760.1 | 690 | 430 | 76.279 | Hu et al., 2016 |
| g76564.t1 | *Preprosomatostatin I | AJB44559.1 | 218 | 114 | 96.491 | Zhenzhen et al., 2014 |
| g79784.t1 | *Preprosomatostatin II | AJB44560.1 | 125 | 94 | 73.404 | Zhenzhen et al., 2014 |
| g56581.t1 | *Preprosomatostatin III | AJB44561.1 | 221 | 12 | 95.833 | Zhenzhen et al., 2014 |
| g42530.t1 | *Prolactin | CAA31060.1 | 382 | 210 | 95.714 | Ma et al., 2016 |
| g75666.t1 | *Prolactin receptor | QIB98245.1 | 1089 | 611 | 86.579 | Ma et al., 2016 |
| g7570.t1 | *60S ribosomal protein L10 | XP_018921966.1 | 439 | 214 | 98.131 | Damzamn et al., 2020 |
| g29499.t1 | *Antithrombin-III | XP_018920986.1 | 845 | 450 | 92.444 | Dam et al 2020 |
| g81491.t1 | *SMAD family member 3 | ABI94729.1 | 877 | 423 | 99.527 | Lin et al., 2019 |
| g94343.t1 | *Secreted Protein Acidic And Cysteine Rich | XP_003447656.1 | 519 | 300 | 83.667 | Lin et al., 2019 |
| g17664.t1 | *Squalene monooxygenase | NP_001103509.1 | 970 | 528 | 89.394 | Ma et al., 2016 |
| g27484.t1 | *Somatostatin Receptor type 1-like | XP_018943223.1 | 749 | 367 | 98.365 | Ma et al., 2016 |
| g16955.t1 | *Somatostatin Receptor type 2-like | XP_018946514.1 | 597 | 337 | 87.834 | Zhenzhen et al., 2014 |
| g82792.t1 | *Somatostatin Receptor type 3-like | XP_018963641.1 | 514 | 261 | 98.084 | Zhenzhen et al., 2014 |
| g12909.t1 | *Somatostatin Receptor type 5-like | XP_018961210.1 | 661 | 443 | 81.49 | Zhenzhen et al., 2014 |
| g32515.t1 | *Signal transducer and activator of transcription 1b | NP_956385.2 | 1161 | 695 | 79.712 | Zhenzhen et al., 2014 |
| g1399.t1 | *Signal transducer and activator of transcription 2 | NP_001258730.1 | 1359 | 852 | 78.873 | Zhenzhen et al., 2014 |
| g23419.t1 | *Signal transducer and activator of transcription 3 | BAH47263.1 | 1675 | 806 | 99.256 | Zhenzhen et al., 2014 |
| g28531.t1 | *Signal transducer and activator of transcription 4 | NP_001004510.1 | 1324 | 679 | 95.582 | Zhenzhen et al., 2014 |
| g48083.t1 | *Signal transducer and activator of transcription 5 | BAH47264.1 | 1549 | 768 | 97.266 | Zhenzhen et al., 2014 |
| g4785.t1 | *Ubiquitin carboxyl-terminal hydrolase 38 | XP_003449754.1 | 1415 | 1013 | 70.582 | Lin et al., 2019 |

**Figure 11.**
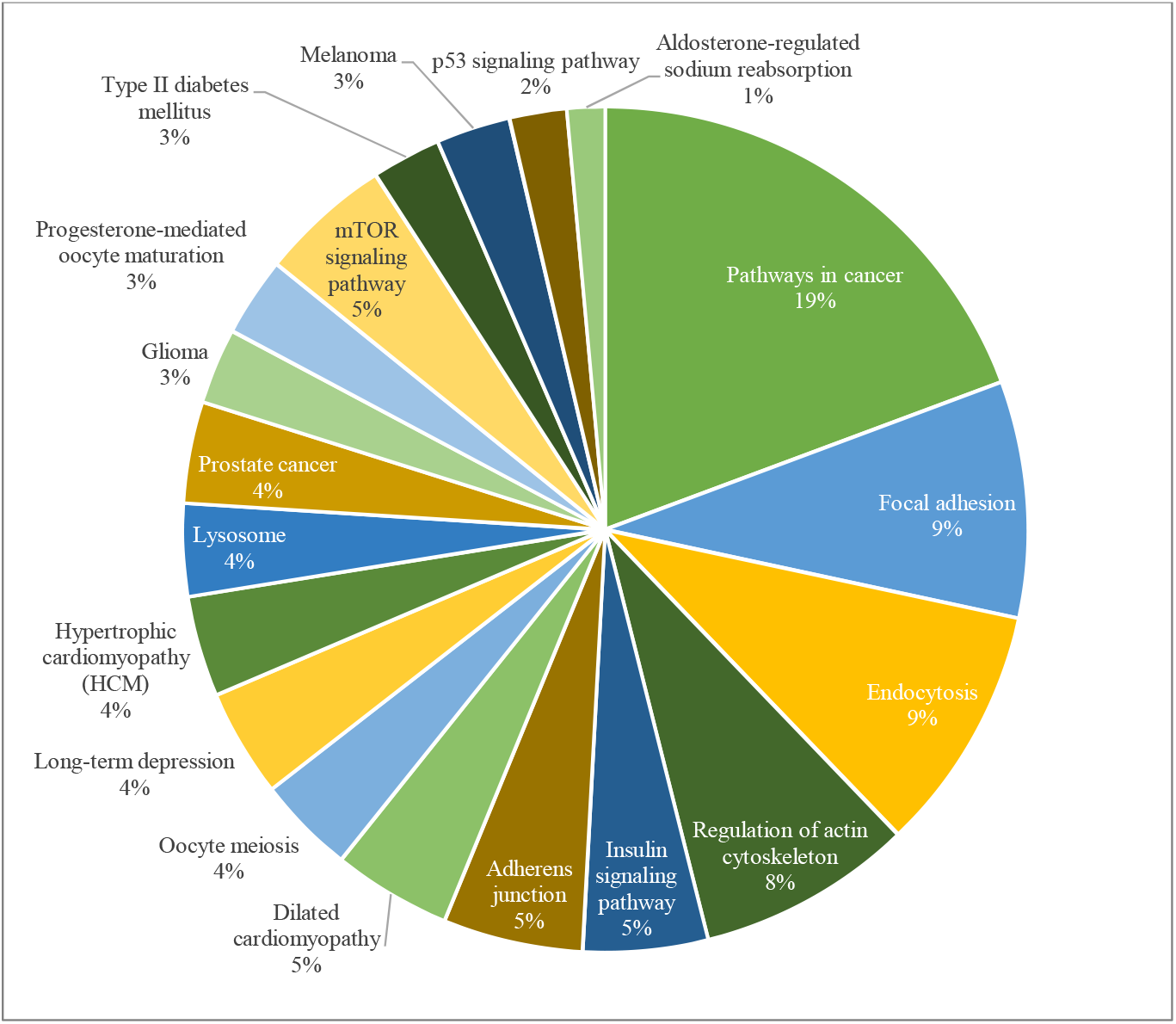
Annotation to growth-related KEGG pathway.

Thus, it can be said that a significant similarity had been exhibited with previously published immune-related genes. For instance, growth hormone (GH) and insulin-like growth factor-binding protein (IGFBP) which are responsible for the regulation of GH and IGF (Zhenzhen et al., 2014) were detected in high similarity in the genome of *T. tambroides* (Table 6). Somatostatins were found to have an inhibitory role in promoting the release of GH (Li & Lin, 2010).

It was found out that regulation of appetite, protein and lipid metabolism, weight gain and muscle growth are also part of the complex growth process (Zhenzhen et al., 2014). A few genes related to appetite and muscle were detected in the genome of *T. tambroides*, for instance, neuropeptide Y (*NPY*), pro-opiomelanocortin (*POMC*), leptin and myostatin (Table 6). These genes play various roles where *NPY* and *POMC* are exerting antagonistic roles in stimulating or inhibiting feeding (Zhenzhen et al., 2014). In addition, myostatin is a negative regulator of muscle growth and its polymorphism is associated with growth traits (Nadjar-Boger & Funkenstein, 2011). In addition, leptin is responsible for the regulation of energy intake and usage (Zhenzhen et al., 2014).

Moroever, genes that encode for proteolytic digestive enzymes (chymotrypsin-like elastase, *cela*) and are related to protein metabolism (cathepsin L, *clsL* and cathepsin K, *clsK*) were detected in the genome of *T. tambroides* as well. In addition, a number of lipid metabolism regulation genes were found in the genome of *T. tambroides*, including lipase C (*lipC*), phospholipase A2 (*pla2*), elongation of very long chain fatty acid family 6 (*elovl6*), apolipoprotein B (*apob*), acetyl-CoA carboxylase (*ACACA, ACACB*) and fatty acid synthase (*FASN*). This can be due the fact that *T. tambroides* is a semi-fatty fish and it contains 4.6-5.2% of muscle crude lipid (Özogul and Özogul, 2007), indicating the importance of lipid metabolism in this fish. Lipase is a key enzyme involved in lipid hydrolysis while apolipoprotein is a lipid-associated protein that regulates lipid homeostasis through the transport of triacylglycerol and phospholipid from the liver to other tissues (Infante & Cahu, 2007). *ACACB* was found not only to be associated with fat yield and percentage, but it also plays role in protein yield as well (Han et al., 2018). *FASN* is an important element in lipid metabolism and its expression could vary due to fatty acid content in both fat and meat (Renaville et al., 2018). Furthermore, glucose-6-phosphatase (*g6pc*) which regulates carbohydrate metabolism were detected in *T. tambroides* as well. These growth-related genes may serve as the possible molecular growth-related markers for further marker-assisted breeding. Further studies are required to confirm the roles of these genes in the growth of *T. tambroides*.

### Orthologs and Phylogenetic Inferences

The phylogenetic relationship of *T. tambroides* with other ray-finned fishes was inferred through the BUSCO supermatrix approach through single-copy orthologs. The BUSCO completeness of each Actinopterygii species was summarized in Table 7. MUSCLE was used to align all the genomic sequences of all 53 species (Edgar, 2004). The phylogenetic tree was plotted using the maximum-likelihood model (ML; IQ-TREE) based on the single-copy and multi-copy orthologs (Figure 12). All the species fall under class Actinopterygii but belong to 18 different orders, including Cypriniformes, Perciformes, Clupeiformes, Cichliformes, Characiformes, Cyprinodontiformes, Anabantiformes, Salmoniformes, Esociformes, Gadiformes, Pleuuronectiformes, Lepisosteiformes, Atheriniformes, Beloniformes, Osteoglossiformes, Batrachoidiformes with Coelacanthiformes and Acipenseriformes rooted as outgroup. As a member of the family *Cyprinidae, T. tambroides* formed a monophyletic cluster with the species within order Cypriniformes, namely species from genus *Sinocyclocheilus, Carassius auratus, Pimephales promelas* and *Danio rerio*.

**Table 7.**
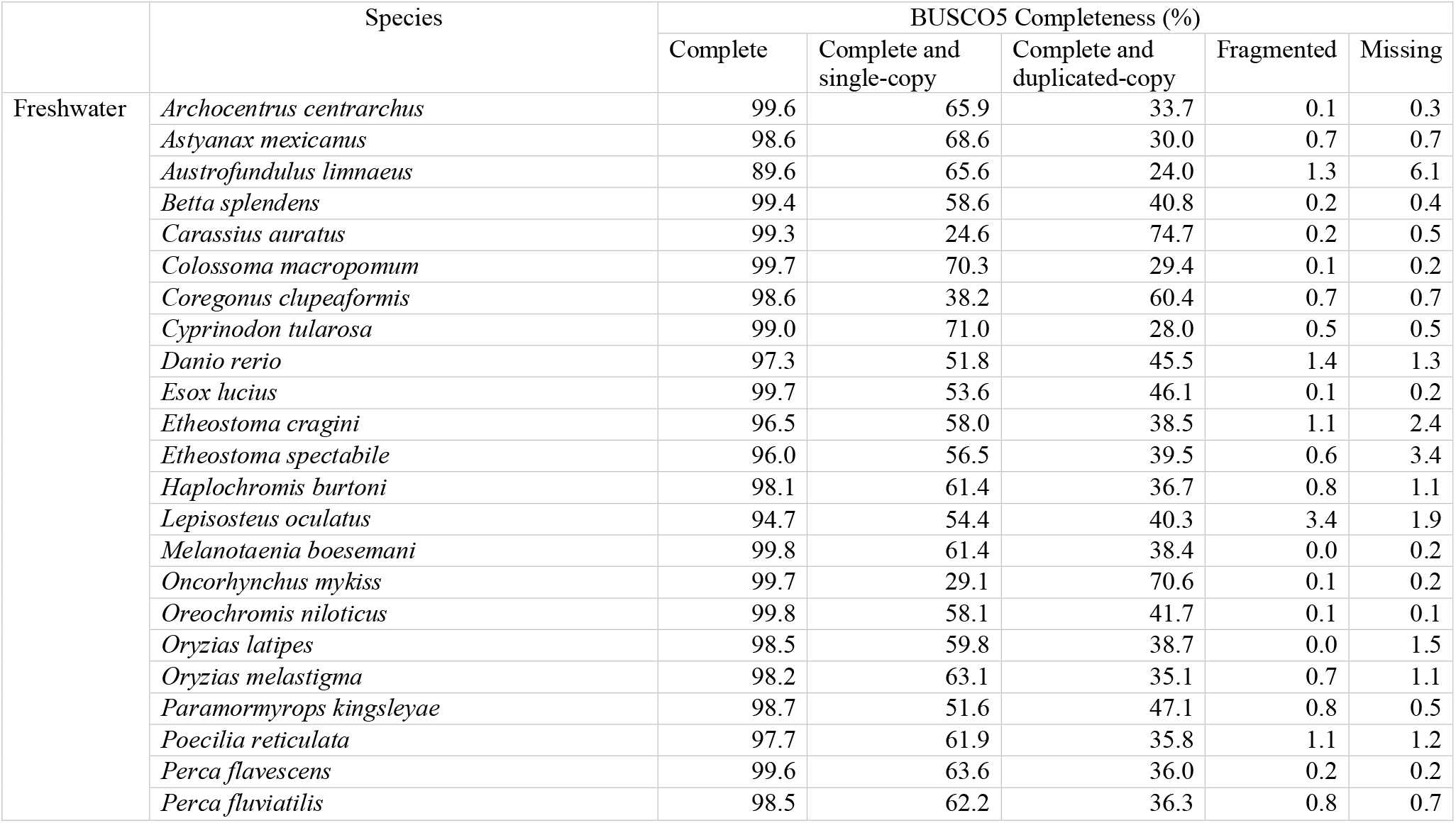
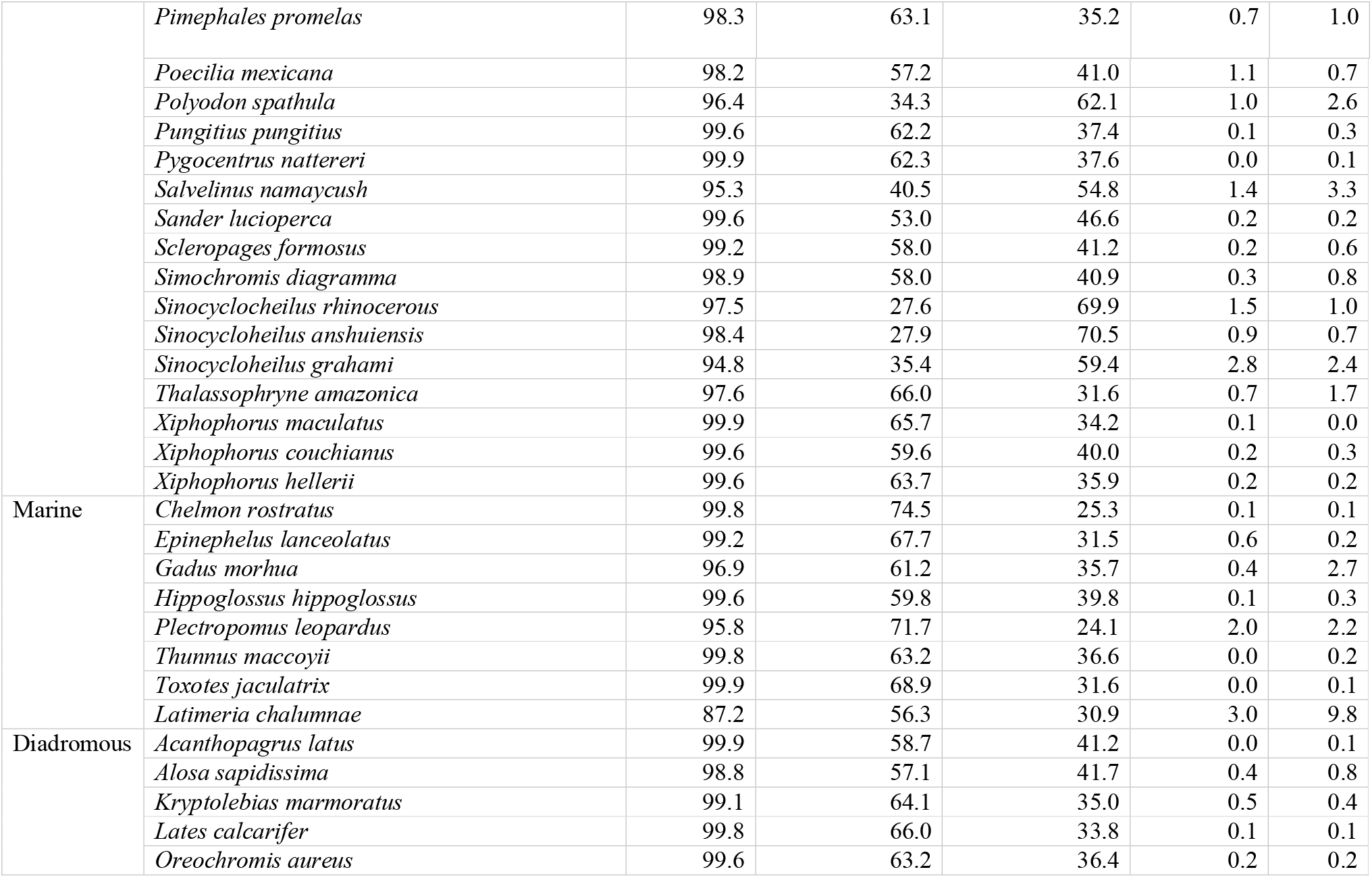
BUSCO completeness of 52 ray-finned fishes for the inference of phylogenetic relationship with *T. tambroides* using BUSCO supermatrix approach.

|  | Species | BUSCO5 Completeness (%) |  |  |  |  |
| --- | --- | --- | --- | --- | --- | --- |
|  |  | Complete | Complete and single-copy | Complete and duplicated-copy | Fragmented | Missing |
| Freshwater | *Archocentrus centrarchus* | 99.6 | 65.9 | 33.7 | 0.1 | 0.3 |
|  | *Astyanax mexicanus* | 98.6 | 68.6 | 30.0 | 0.7 | 0.7 |
|  | *Austrofundulus limnaeus* | 89.6 | 65.6 | 24.0 | 1.3 | 6.1 |
|  | *Betta splendens* | 99.4 | 58.6 | 40.8 | 0.2 | 0.4 |
|  | *Carassius auratus* | 99.3 | 24.6 | 74.7 | 0.2 | 0.5 |
|  | *Colossoma macropomum* | 99.7 | 70.3 | 29.4 | 0.1 | 0.2 |
|  | *Coregonus clupeaformis* | 98.6 | 38.2 | 60.4 | 0.7 | 0.7 |
|  | *Cyprinodon tularosa* | 99.0 | 71.0 | 28.0 | 0.5 | 0.5 |
|  | *Danio rerio* | 97.3 | 51.8 | 45.5 | 1.4 | 1.3 |
|  | *Esox lucius* | 99.7 | 53.6 | 46.1 | 0.1 | 0.2 |
|  | *Etheostoma cragini* | 96.5 | 58.0 | 38.5 | 1.1 | 2.4 |
|  | *Etheostoma spectabile* | 96.0 | 56.5 | 39.5 | 0.6 | 3.4 |
|  | *Haplochromis burtoni* | 98.1 | 61.4 | 36.7 | 0.8 | 1.1 |
|  | *Lepisosteus oculatus* | 94.7 | 54.4 | 40.3 | 3.4 | 1.9 |
|  | *Melanotaenia boesemani* | 99.8 | 61.4 | 38.4 | 0.0 | 0.2 |
|  | *Oncorhynchus mykiss* | 99.7 | 29.1 | 70.6 | 0.1 | 0.2 |
|  | *Oreochromis niloticus* | 99.8 | 58.1 | 41.7 | 0.1 | 0.1 |
|  | *Oryzias latipes* | 98.5 | 59.8 | 38.7 | 0.0 | 1.5 |
|  | *Oryzias melastigma* | 98.2 | 63.1 | 35.1 | 0.7 | 1.1 |
|  | *Paramormyrops kingsleyae* | 98.7 | 51.6 | 47.1 | 0.8 | 0.5 |
|  | *Poecilia reticulata* | 97.7 | 61.9 | 35.8 | 1.1 | 1.2 |
|  | *Perca flavescens* | 99.6 | 63.6 | 36.0 | 0.2 | 0.2 |
|  | *Perca fluviatilis* | 98.5 | 62.2 | 36.3 | 0.8 | 0.7 |
|  | *Pimephales promelas* | 98.3 | 63.1 | 35.2 | 0.7 | 1.0 |
|  | *Poecilia mexicana* | 98.2 | 57.2 | 41.0 | 1.1 | 0.7 |
|  | *Polyodon spathula* | 96.4 | 34.3 | 62.1 | 1.0 | 2.6 |
|  | *Pungitius pungitius* | 99.6 | 62.2 | 37.4 | 0.1 | 0.3 |
|  | *Pygocentrus nattereri* | 99.9 | 62.3 | 37.6 | 0.0 | 0.1 |
|  | *Salvelinus namaycush* | 95.3 | 40.5 | 54.8 | 1.4 | 3.3 |
|  | *Sander lucioperca* | 99.6 | 53.0 | 46.6 | 0.2 | 0.2 |
|  | *Scleropages formosus* | 99.2 | 58.0 | 41.2 | 0.2 | 0.6 |
|  | *Simochromis diagramma* | 98.9 | 58.0 | 40.9 | 0.3 | 0.8 |
|  | *Sinocyclocheilus rhinoceros* | 97.5 | 27.6 | 69.9 | 1.5 | 1.0 |
|  | *Sinocyclocheilus anshuiensis* | 98.4 | 27.9 | 70.5 | 0.9 | 0.7 |
|  | *Sinocyclocheilus grahami* | 94.8 | 35.4 | 59.4 | 2.8 | 2.4 |
|  | *Thalassophryne amazonica* | 97.6 | 66.0 | 31.6 | 0.7 | 1.7 |
|  | *Xiphophorus maculatus* | 99.9 | 65.7 | 34.2 | 0.1 | 0.0 |
|  | *Xiphophorus couchianus* | 99.6 | 59.6 | 40.0 | 0.2 | 0.3 |
|  | *Xiphophorus hellerii* | 99.6 | 63.7 | 35.9 | 0.2 | 0.2 |
| Marine | *Chelmon rostratus* | 99.8 | 74.5 | 25.3 | 0.1 | 0.1 |
|  | *Epinephelus lanceolatus* | 99.2 | 67.7 | 31.5 | 0.6 | 0.2 |
|  | *Gadus morhua* | 96.9 | 61.2 | 35.7 | 0.4 | 2.7 |
|  | *Hippoglossus hippoglossus* | 99.6 | 59.8 | 39.8 | 0.1 | 0.3 |
|  | *Plectropomus leopardus* | 95.8 | 71.7 | 24.1 | 2.0 | 2.2 |
|  | *Thunnus maccoyii* | 99.8 | 63.2 | 36.6 | 0.0 | 0.2 |
|  | *Toxotes jaculatrix* | 99.9 | 68.9 | 31.6 | 0.0 | 0.1 |
|  | *Latimeria chalumnae* | 87.2 | 56.3 | 30.9 | 3.0 | 9.8 |
| Diadromous | *Acanthopagrus latus* | 99.9 | 58.7 | 41.2 | 0.0 | 0.1 |
|  | *Alosa sapidissima* | 98.8 | 57.1 | 41.7 | 0.4 | 0.8 |
|  | *Kryptolebias marmoratus* | 99.1 | 64.1 | 35.0 | 0.5 | 0.4 |
|  | *Lates calcarifer* | 99.8 | 66.0 | 33.8 | 0.1 | 0.1 |
|  | *Oreochromis aureus* | 99.6 | 63.2 | 36.4 | 0.2 | 0.2 |

**Figure 12.**
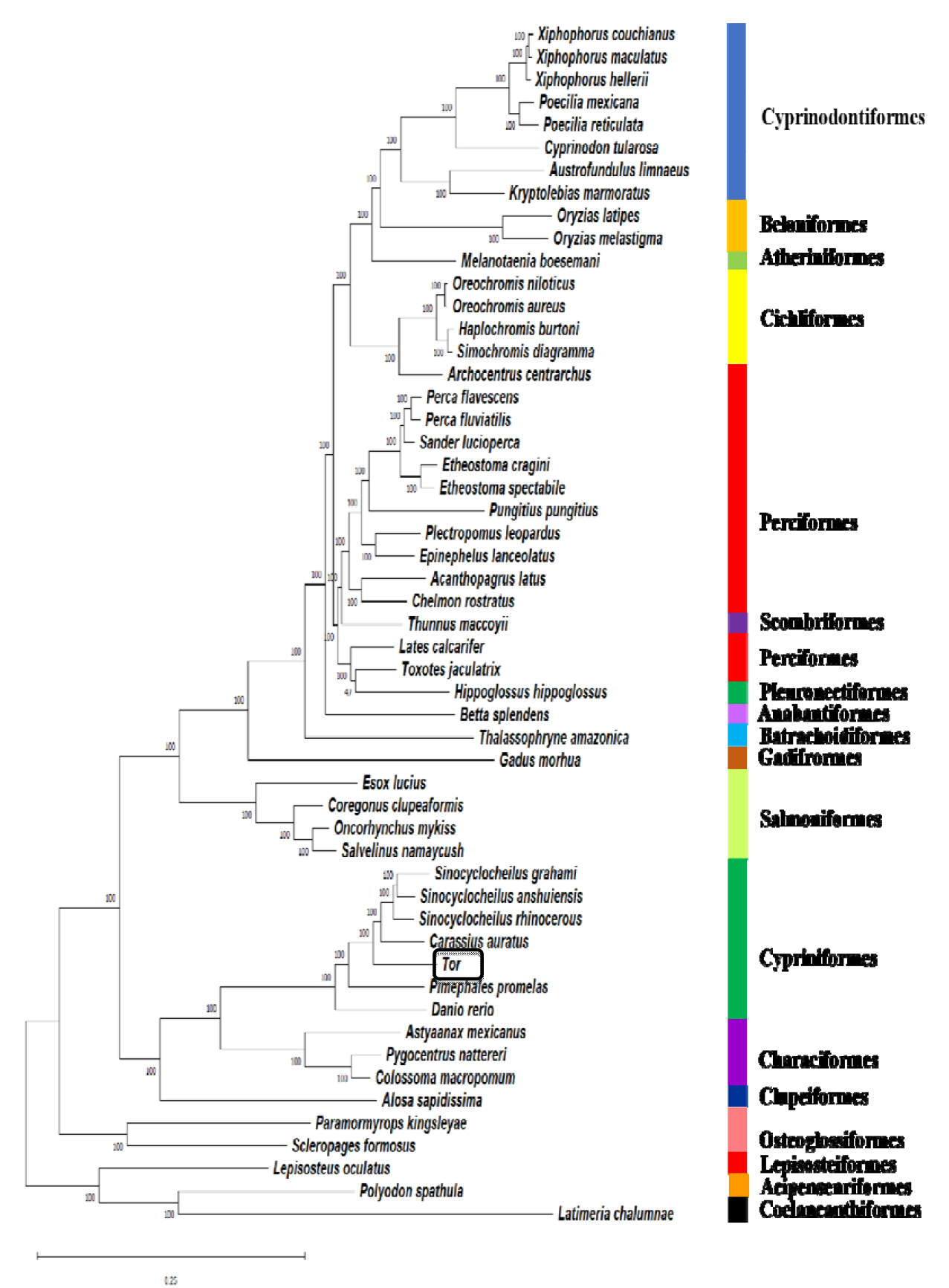
Maximum-likelihood tree plotted on 53 Actinopterygii species inferred from a supermatrix of 3,640 BUSCO. The species were classified based on their respective orders.

## Conclusion

We present the first Malaysian mahseer (*T. tambroides*) genome assembled with low-coverage Nanopore long reads and high-coverage Illumina short reads. *De novo* genomic assembly had generated a draft genome with an estimated genome size of 1.5 Gb [84.3% BUSCO completeness (Actinopterygii_odb10)]. 392,346 SSRs had been identified from the genome with dinucleotide repeats AT/TA as the most common SSR. A total of 96,736 non-redundant protein-coding sequences was predicted and used for later functional annotation. Predicted protein sequences had mapped to 304 known KEGG pathways with signal transduction as the highest representation. Furthermore, genes showing significant similarity to published growth- and immune-related genes were identified, hoping to serve as a potential marker for future molecular breeding of *T. tambroides*. In addition, the first genome-based evolutionary relationship of *T. tambroides* between other ray-finned fishes had been inferred using a Maximum-likelihood tree. It is hoped that this genomic data of *T. tambroides* could be a more powerful tool that continues to resolve questions of species identification, evolutionary biology, morphological variations, sequences related to sex differentiation, growth, reproduction, and immune, which are helpful for further conservation of *Tor* species.

## CrediT author statement

**Lau Melinda Mei Lin:** Writing-Original Draft, Data curation, Conceptualization. **Lim Leonard Whye Kit:** Data Curation, Writing-Original Draft, Conceptualization. **Chung Hung Hui:** Conceptualization, Funding acquisition, Writing-Review and Editing. **Han Ming Gan:** Methodology, Conceptualization, Writing-Review and Editing.

## Acknowledgements

This work was fully funded by Sarawak Research and Development Council through the Research Initiation Grant Scheme with grant number RDCRG/RIF/2019/13 awarded to H. H. Chung.

